# Small molecules targeting *BRCA1*-deficient cancer cells

**DOI:** 10.64898/2026.10.21.760761

**Authors:** Arunasalam Navaraj, Shengliang Zhang, Chris Egg-Krings, Kevin Si, Jean-Nicolas Gallant, Jay Gopal, Sriya S.P. Datla, David T. Dicker, Dinara Ryspayeva, Wafik S. El-Deiry

## Abstract

*BRCA1* mutation-carriers frequently develop triple-negative breast cancers that are resistant to cancer therapies. We used our established luciferase-expressing *BRCA1*-deficient human mammary epithelial-derived cell line series with stable shRNA-mediated *BRCA1*-knockdown, H-RasG12V-mediated transformation, and *TP53*-deficiency (PMID: 20038817) to perform a high-throughput-screen using the National Cancer Institute (NCI) chemical library Diversity Set II of 1,440 small molecules and noninvasive bioluminescence imaging. We selected five (NSC667251, NSC636948, NSC21683, NSC281383, and NSC136513) compounds that reproducibly induced cell death in engineered *BRCA1*-knockdown cells and in *BRCA1*-deficient human cancer cell lines. In p53-mutant MDA-MB-231 and SKBr3 lines, the compounds induced apoptosis and PARP cleavage by 12 hours. Sub-G1 flow cytometric analysis demonstrated maximal apoptosis induction by 48 hours of drug exposure. p53 protein stabilization and phosphorylation were observed with the selected compounds in *BRCA1*-deficient wild-type *TP53*-expressing cells. Drug-treated *BRCA1*-knockdown MCF10A cells and *BRCA1*-mutant HCC-1937 breast cancer cells showed γ-H2AX foci and phospho-H2AX by immunoblotting indicating genotoxicity and DNA damage as contributing factors for apoptosis. NSC636948 was previously characterized as an inhibitory ligand of peroxisome proliferator-activated β/δ (PPARβ/δ) and both NSC636948 and NSC667251 act as inverse agonists of PPARβ/δ. Kaplan–Meier survival analysis of a combined breast cancer cohort (n = 1,959) derived from the METABRIC, demonstrated that patients with high *PPARD* expression exhibited significantly worse overall survival compared with those with low *PPARD* expression (log-rank p = 0.008). The selected compounds provide candidate therapeutic leads as well as insights into molecular targets for further evaluation in *BRCA1*-deficient tumor models and for assessment in combination therapy strategies.

## Introduction

Breast cancer is the most frequently diagnosed malignancy in women worldwide. GLOBOCAN estimates place the 2022 global burden at approximately 2.3 million new cases and 666,000 deaths, making it the leading cause of cancer death among women in most countries surveyed [1]. The latest GLOBOCAN statistics, published in July 2026, describe a global cancer incidence estimated at 20.6 million in 2024 and this includes a breast cancer incidence of 2.4 million new cases as the second most commonly diagnosed cancer worldwide (11.8% of all new cancer cases), and with 694,000 deaths (7.4% of all cancer deaths worldwide) (https://acsjournals.onlinelibrary.wiley.com/doi/10.3322/caac.70090).

Most hereditary breast cancer is attributable to germ-line mutation of *BRCA1*, the breast and ovarian cancer susceptibility gene mapped to chromosome 17q21 and cloned in 1994 [2,3]. Prospective cohort analysis of nearly 10,000 carriers established a cumulative breast cancer risk of 72% by age 80 in women carrying a pathogenic *BRCA1* variant, with risk rising significantly from the third decade of life [4].

*BRCA1* encodes a large, multifunctional tumor suppressor that participates in transcriptional regulation, cell cycle checkpoint control, ubiquitin ligase activity, and centrosome duplication. The most consequential tumor suppressive activity of *BRCA1*, however, is in the maintenance of genomic integrity where it serves a role in mediating homologous recombination (HR) in the repair of DNA double-strand breaks [5,6] and for protection of stalled replication forks from nucleolytic degradation [7,8]. Mammalian cells lacking functional *BRCA1* accumulate chromosomal aberrations, fail to condense metaphase chromosomes normally, acquire supernumerary centrosomes, and display marked hypersensitivity to ionizing radiation and to DNA-damaging chemotherapeutic agents [9,10,11]. This hypersensitivity phenotype is the biological foundation of every therapeutic strategy currently directed at *BRCA1*-deficient disease, taking advantage of a synthetic lethality for example with poly-ADP ribose polymerase (PARP) inhibition.

Clinically, tumors arising in *BRCA1* mutation carriers are overwhelmingly of basal-like transcriptional subtype and are, in most series, triple-negative — lacking expression of estrogen receptor, progesterone receptor, and HER2 [12,13,14]. *BRCA1*-deficient tumors are typically ineligible for endocrine therapy or for HER2-directed therapeutic agents, and cytotoxic chemotherapy remains the backbone of their management, along with PARP inhibitors. It should be noted that triple-negative breast cancer (TNBC) and breast cancers with *BRCA1*-loss are overlapping rather than equivalent categories: TNBC is transcriptionally heterogeneous, comprising several distinct subtypes of which only a subset carries *BRCA1* alteration [15].

A second and frequently underappreciated feature of these tumors is the near-obligate co-occurrence of *TP53* mutation. In *BRCA1*-associated basal-like carcinomas, sequencing combined with functional allele analysis has detected *TP53* mutation in 97% of cases, with a distinctive enrichment for complex, protein-truncating alterations relative to sporadic tumors [16,17]; large-scale molecular profiling of the basal-like subtype has returned comparable frequencies [18]. Loss of p53-checkpoint function is permissive for the survival of cells carrying the genomic instability that BRCA1 loss produces, but it may confer resistance to chemotherapy and radiation. Because p53 is the major component of the DNA Damage Response (DDR) that ultimately leads to cellular apoptosis, its loss blunts the death response that genotoxic therapy is intended to trigger and can do so in tumors with HR repair defect that should make them vulnerable. A therapeutic agent capable of killing *BRCA1*-deficient cells through a route that does not require wild-type p53 would therefore address a specific and well-defined gap, yet p53 status is rarely incorporated into screening design as a selection criterion.

The prevailing therapeutic strategy for *BRCA1*-deficient tumors exploits synthetic lethality. Two groups demonstrated in 2005 that inhibition PARP selectively eliminates *BRCA1*- and *BRCA2*-deficient cells while sparing HR-proficient cells [19,20], and this observation has since produced approved agents with demonstrable clinical benefit. Olaparib and talazoparib each prolonged progression-free survival (PFS) relative to physician’s-choice chemotherapy in germline *BRCA1/2*-mutant, HER2-negative metastatic disease [21,22], and adjuvant olaparib improved invasive disease-free and overall survival (OS) in high-risk early breast cancer [23]. In the metastatic setting, the PFS advantage of olaparib did not translate into a significant OS benefit at the final analysis (median 19.3 versus 17.1 months) [24]. Acquired resistance is near-universal and arises through multiple partly independent routes. These include loss of *53BP1* or of the downstream shieldin complex, which permits DNA end resection and partial restoration of HR in a *BRCA1*-null background [25,26], secondary reversion mutations that restore the *BRCA1* reading frame [27], re-establishment of replication fork protection, and increased drug efflux [28,29]. Platinum chemotherapy exploits the same repair defect and is more active than taxane in germline *BRCA1/2*-mutant TNBC with response rates of 68% versus 33% in the randomized TNT trial, but responses were not durable and the agent is not selective for tumor tissue [30]. Agents acting through mechanisms orthogonal to PARP trapping, and retaining activity where HR has been partially restored, remain an unmet need.

We reasoned that drug discovery targeting *BRCA1*-deficient tumors may be approached phenotypically than by target-directed design. Retrospective analysis of first-in-class small molecules approved between 1999 and 2008 found that phenotypic screening outperformed target-based approaches as a source of genuinely novel mechanisms, largely because it imposes no prior hypothesis about which protein must be engaged [31].

We sought to establish an isogenic model system for the screening where background *BRCA1*-deficiency is genetically defined and consistent across drug-screened comparator cell lines. Breast cancer cell lines carry extensive and heterogeneous genomic alterations, so differential sensitivity cannot be easily attributed to *BRCA1*-loss per se. We therefore employed a panel of isogenic derivatives of the immortalized but non-tumorigenic human mammary epithelial line MCF10A [32], in which *BRCA1* and *TP53* were stably depleted alone or in combination, with or without oncogenic H-Ras, alongside a constitutive luciferase reporter [33]. The *BRCA1*-deficient transformed human mammary epithelium-derived cell lines reproduced hallmark features of BRCA1-deficient tumor biology, i.e. chromosome condensation defects, centrosome amplification, genomic instability, reduced apoptosis, increased invasiveness, and tumorigenicity in the Ras-transformed derivatives, while permitting direct comparison against a *BRCA1*-proficient control differing at a single locus [33]. We designed the model *BRCA1*-deficient drug screening such that the viability readout can be longitudinal. The luciferase reporter allowed the same wells to be imaged repeatedly by bioluminescence over several days rather than sampled once at the endpoint, which preserved the detection of compounds with delayed cytotoxic kinetics that a single-time-point assay might not detect.

We applied our established *BRCA1*-deficient human mammary epithelial cell-derived platform to the complete set of 1,440 compounds of the National Cancer Institute Developmental Therapeutics Program Diversity Set. The rationale for screening the set in its entirety, rather than a preselected subset, rests on how it was assembled. Each member was drawn from the NCI open chemical repository of roughly 140,000 compounds and selected computationally so that the collection maximizes pharmacophore coverage while favoring conformationally rigid scaffolds, with the consequence that a library of this modest size samples a disproportionately wide region of chemical space [34]. The compounds screened were structurally unrelated to the oncology pharmacopeia, are indexed by permanent NSC accession numbers, and are freely available in resupply quantities for mechanistic follow-up, all features that made the set well suited to an unbiased, target-agnostic search and to the practical demands of hit validation. Screening the chemical library at a fixed concentration maximized the probability of encountering scaffolds with no established anti-cancer precedent, at a cost of an assay burden that longitudinal bioluminescence imaging made tractable.

We hypothesized that *BRCA1*-loss generates a cell-intrinsic vulnerability that can be engaged pharmacologically by agents that may act independently of PARP-inhibition and independently of *TP53*, and that such agents could be recovered from an unbiased chemical library by a selectivity-based phenotypic screen.

We report the identification of compounds with selective cytotoxicity towards *BRCA1*-deficient transformed human mammary epithelial cells or established tumor cell lines with *BRCA1*-defects, as well as compounds that display cytotoxicity towards *BRCA1*-deficient tumor cells regardless of mammary gland cellular origin. We characterize the isogenic model system, describe the primary screen and the counter-screening cascade used to eliminate nonspecifically cytotoxic agents, establish that the surviving compounds that induce apoptosis in *BRCA1*-depleted mammary epithelial cells and in *BRCA1*-mutant and *BRCA1*-null breast cancer lines while sparing normal fibroblasts and normal mammary epithelium. We preliminarily investigate the mechanism of cell death through γ-H2AX focus formation, PARP cleavage, and p53 stabilization. Together these data establish six compounds as genotoxic agents that remain effective in the *TP53*-deficient background in which *BRCA1*-associated breast cancer characteristically arises and additional compounds that display tumor cell-specific cytotoxicity associated with *BRCA1*-deficiency regardless of the mammary gland cellular origin of the tumor cells.

## Methods

### Cell lines

The cell lines used throughout this study were generated and characterized previously in our laboratory and are described in detail elsewhere [33]. Immortalized human mammary epithelial MCF10A cells were first transduced with a firefly luciferase reporter (ML), then subjected to stable, retrovirally delivered shRNA knockdown of *BRCA1* and/or *TP53* with FACS-based selection for the linked fluorescent markers and finally transduced with mutant H-Ras (H-Ras V12) to generate transformed derivatives. All cell lines used in this study derive from immortalized human mammary epithelial MCF10A cells stably expressing a firefly luciferase reporter construct, designated ML. Line names record the modifications each derivative carries, with B denoting shRNA-mediated knockdown of *BRCA1*, P denoting shRNA-mediated knockdown of p53, and R denoting expression of activated H-Ras. On this scheme, MLB and MLR carry BRCA1 knockdown or Ras transformation alone; MLBP combines knockdown of both *BRCA1* and *TP53*; MLBR and MLPR pair Ras transformation with knockdown of *BRCA1* or of *TP53* respectively; and MLBPR carries all three alterations.

### Cell viability

The viability of cells was measured using an ATP-based luminescence test (CellTiter-Glo viability assay; Promega) carried out 96 hours after the treatment with compounds.

### Flow cytometry

The amount of apoptosis was determined based on the evaluation of sub-G1 DNA content via flow cytometry. The collected cells were stained with propidium iodide after fixing in ethanol.

### Immunofluorescence

DNA damage was tested based on the presence of gamma-H2AX protein expression by immunofluorescence at time-points indicated in the figure legends. DAPI was used for nuclear counterstaining.

### Western blots

Whole-cell lysates were subjected to Western blot analysis to detect the amount of gamma-H2AX, phosphorylated (Ser 15) and total amounts of the p53 protein (antibody from Cell signaling), and cleaved poly(ADP-ribose) polymerase. All the experiments were performed at least three times independently. The following antibodies were used:

1. Phospho H2AX Antibody - Catalog # 05-636, Sigma Aldrich, Clone JBW301
2. p53 Antibody - Catalog # SC-126 HRP, Santa Cruz
3. Phospho p53 (Serine 15) Antibody - Catalog # 9286, Cell Signaling
4. PARP Antibody - Catalog # 9542, Cell Signaling
5. Ran Antibody - Catalog # 610341, BD Transduction Laboratories

### Secondary compound activity screening

Thirteen compounds (NSC 300289, 87084, 367089, 13051, 93427, 136513, 667251, 281383, 275971, 636948, 21683, 98938, and 36758) were evaluated against the NCI-60 human tumor cell line panel, comprising 60 cell lines derived from nine tissues of origin (breast, central nervous system, colon, leukemia, melanoma, lung, ovarian, prostate, and renal), maintained by the National Cancer Institute’s Developmental Therapeutics Program. Growth inhibition was assessed using the sulforhodamine B (SRB) protein-binding assay [45], with compound potency for each cell line expressed as GI50 (concentration producing 50% growth inhibition), reported as −log10(GI50).

### Data retrieval and normalization in secondary compound screens

Per-cell-line compound activity data were retrieved from CellMiner (v2.15; discover.nci.nih.gov/cellminer), an NCI-maintained database and analysis suite for the NCI-60 [46]. For each compound, activity values were converted to z-scores across the 60-cell-line panel (mean = 0, SD = 1), reflecting the average across replicate experiments passing CellMiner’s quality-control criteria.

### Secondary screening compound analysis for tumor tissue-level aggregation

Per-cell-line z-scores were averaged within each of the nine NCI-60 tissue subpanels for each compound. Cell lines lacking a validated activity value for a given compound were excluded from that compound’s tissue-level average rather than imputed.

### Core antiproliferative family of screened compounds

The six-compound core antiproliferative family (NSC 136513, 667251, 281383, 275971, 636948, and 21683) was defined by pairwise Pearson correlation of each compound’s NCI-60 activity z-score pattern against the other members of the group, computed in R. All six compounds showed mean within-group correlations of r = 0.53–0.72; no other screened compound exceeded r = 0.5 by the same measure (maximum r = 0.43, NSC 93427).

### Data visualization for screened compound tumor type activity in NCI-60 tissue-of-origin subpanels

Tissue-averaged z-scores were visualized as a heatmap using R (R Core Team) and the ggplot2 package [47]. Cells are colored on a diverging red–white–blue scale centered at zero, with red indicating relative sensitivity and blue indicating relative resistance.

### Statistical Analysis

Data are presented as mean ± standard deviation from at least three independent experiments. Statistical comparisons between groups were performed using two-tailed Student’s t-tests or one-way ANOVA with post hoc multiple-comparison correction where appropriate. P values <0.05 were considered statistically significant.

## Results

### High-throughput functional, bioluminescence imaging-based screen of NCI Diversity Set identifies small molecule chemical compounds with *BRCA1*-selective cytotoxicity

To identify small molecules that preferentially kill cells lacking *BRCA1*, we performed a cell-based, high-throughput functional screen using non-invasive bioluminescence imaging as a surrogate readout for viability. Isogenic MCF10A-luciferase cells, either *BRCA1*-proficient (ML) or stably *BRCA1*-depleted (MLB), were seeded at 10,000 cells per well in white opaque 96-well plates and treated with individual compounds from the National Cancer Institute Developmental Therapeutics Program Diversity Set II. A total of 1,440 compounds were screened at a final concentration of 5 µM, and total photon flux was quantified at 24-hour intervals over 96 hours; vehicle (DMSO)-treated wells on every plate served as internal controls (example in **Figure 1A**). Loss of luciferase signal relative to the paired isogenic control was used as the primary indicator of cell death. Molecular structures of small molecules that appeared to be preferentially cytotoxic to cells with WT *BRCA1* and those that appeared to be preferentially cytotoxic to cells with *BRCA1*-knockdown in the chemical library screen are shown in **Figure 1B** and **1C**, respectively. The chemical library screen identified compounds that fell into three classes, depending on whether they were selective for *BRCA1* loss (**Table 1**).

**Figure 1.**
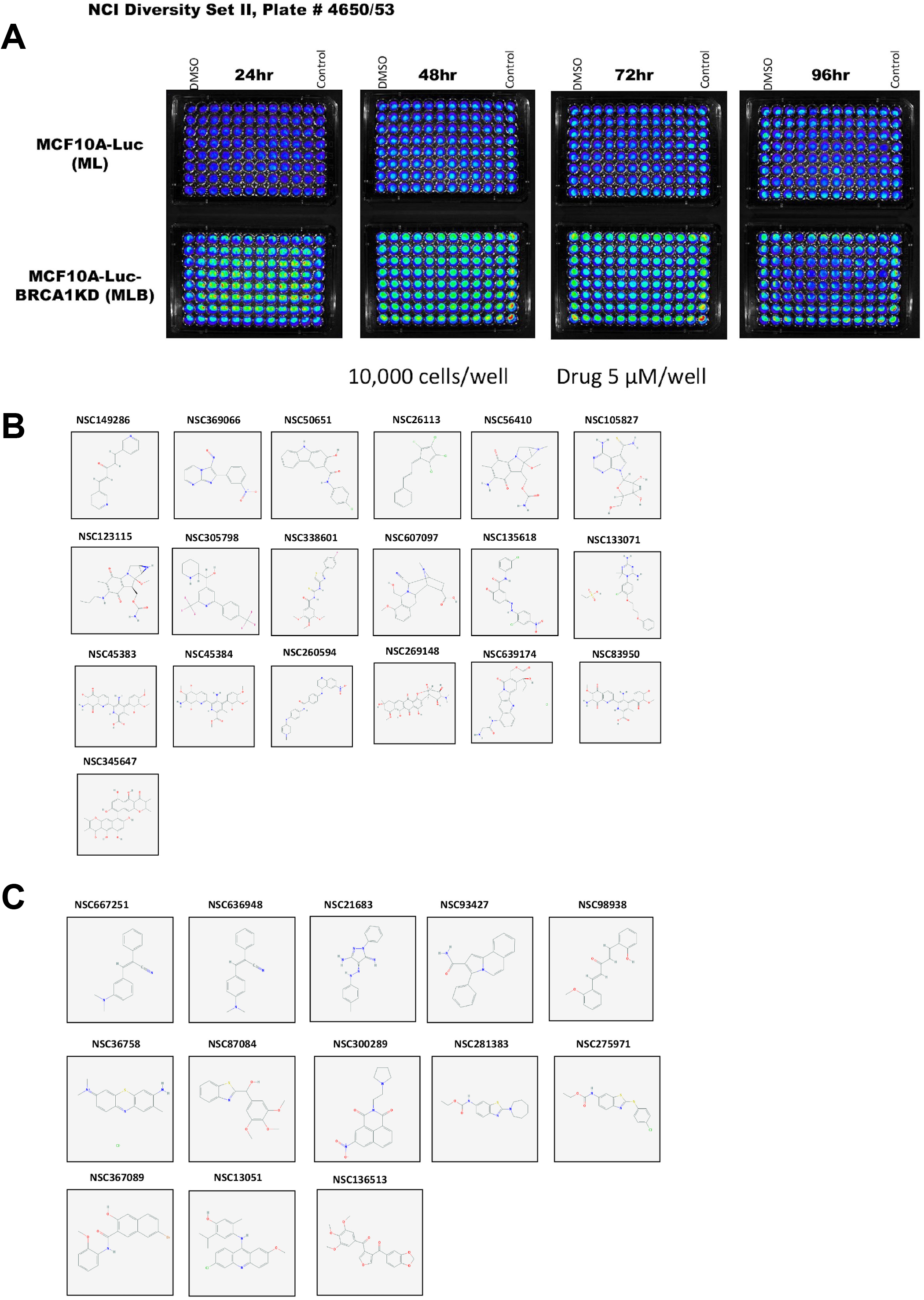
Identification of novel anti-cancer agents through high-throughput cell-based drug screening for BRCA1 pathway specific synthetic lethality. (A) Luciferase expressing isogenic cell lines were screened with the National Cancer Institute (NCI) Developmental Therapeutics Program (DTP) Diversity Set II library of 1440 small molecule compounds. An example is shown with library plate #4650/53 comparing phenotypes of the genetically engineered cell lines MCF10A-Luc (ML) and MCF10A-Luc-BRCA1KD (MLB). Each well of the 96 well plate had 10,000 cells plated and drugs were administered at 5 μM for each compound. A time course is shown for bioluminescence imaging at 24, 48, 72, and 96 hours. (B) Molecular structures of small molecules that appeared to be preferentially cytotoxic to cells with WT BRCA1 in the chemical library screen. (C) Molecular structures of small molecules that appeared to be preferentially cytotoxic to cells with BRCA1-knockdown in the chemical library screen.

**Table 1.** Summary of small molecules identified from the high throughput drug screen.

| Total Compounds | Wild Type Specific | BRCA1 K.D. Specific | Both Common |
| --- | --- | --- | --- |
| 1440 | 19 (1.32%) | 13 (0.90%) | 30 (2.08%) |

Nineteen compounds (1.32% of the library) reduced luciferase signal preferentially in *BRCA1*-proficient ML cells and were classified as wild-type–specific. Thirteen compounds (0.90%) produced a pronounced, time-dependent loss of signal in *BRCA1*-depleted MLB cells while leaving isogenic ML controls largely intact and were designated *BRCA1*-knockdown–specific. A further 30 compounds (2.08%) reduced signal equivalently in both genotypes; because such compounds cannot distinguish *BRCA1* status, they were considered generally cytotoxic and excluded from further analysis (**Table 1**). The overall hit rate across all three classes was 4.3% (62 of 1,440), and the *BRCA1*-selective class — the group of interest for this study — represented fewer than one percent of the library, underscoring the stringency of the selection criteria.

### *BRCA1*-selectivity tracks *BRCA1*-loss independently of *TP53* status and Ras transformation

The 13 BRCA1-knockdown–specific compounds were re-tested across the isogenic MCF10A series at two concentrations, 5 µM and 1 µM, with luciferase signal read at 96 hours (**Figures 2**). Five genotypes were evaluated: parental MCF10A-Luc (ML), *BRCA1*-depleted (MLB), *BRCA1/TP53* double-depleted (MLBP), H-Ras–transformed (MLR), and *BRCA1*-depleted, H-Ras–transformed (MLBR). This design was chosen to separate three variables that are conflated in most established cancer lines — *BRCA1* status, *TP53* status, and oncogenic transformation — against an otherwise identical genetic background.

**Figure 2.**
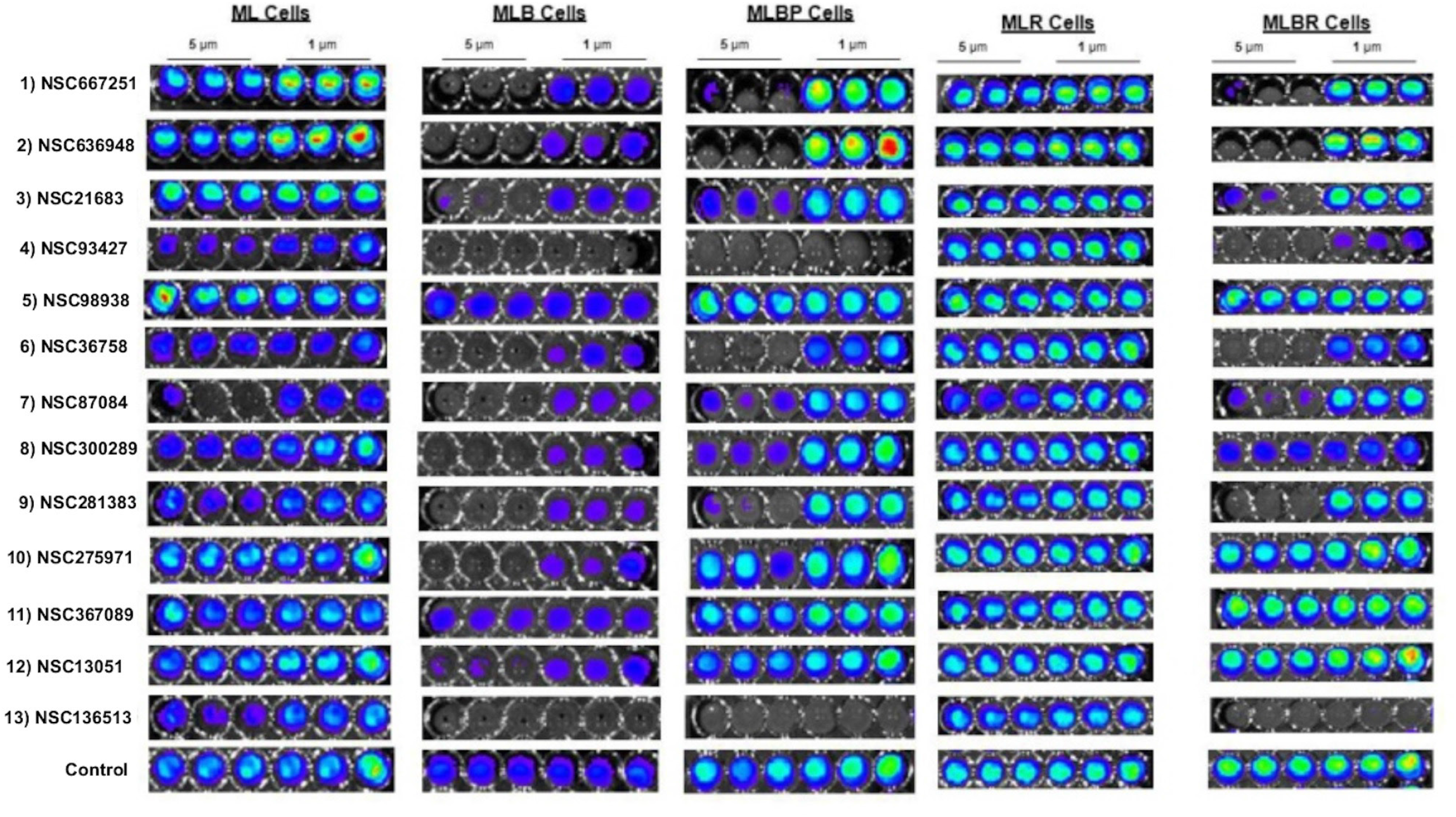
Secondary screening for synthetic lethality by phenotypic loss of cell viability among BRCA1-deficient isogenic series of cell lines derived from human mammary epithelial cells treated with candidate cytotoxic small molecule compounds form primary chemical library screening. The cell lines shown include ML (MCF10A-Luc), MLB (MCF10A-Luc-BRCA1-KD, MLBP (MCF10A-Luc-BRCA1-KD-p53), MLR (MCF10A-Luc-Ras), and MLBR (MCF10A-Luc-BRCA1-KD-Ras). Cell lines were treated with 1 or 5 μM as indicated of the small molecules (NCI DTP Diversity set II NSC numbers shown on the left) and imaged at 96 hours. Black well indicate loss of cells while blue or red wells indicate more cells present. Each experimental condition was tested in triplicate.

In the untransformed pair, compound-induced loss of signal was consistently greater in MLB than in ML cells at both concentrations, and the differential was retained at 1 µM for most hits, indicating that selectivity was not an artifact of a saturating dose (**Figure 2**). The same pattern held after Ras-mediated transformation: MLBR cells were markedly more sensitive than MLR cells at both 5 µM and 1 µM, confirming that *BRCA1* loss, rather than the transformed state itself, governed the response (**Figure 2**). Importantly, MLBP cells, which lack both BRCA1 and p53, were killed as efficiently as MLB cells. Because loss of p53 function is a well-recognized source of chemoresistance and frequently co-occurs with *BRCA1* mutation in human tumors, this p53 independence was regarded as a favorable property and was used as a selection criterion going forward.

Five of the 13 compounds — NSC98938, NSC300289, NSC275971, NSC367089 and NSC13051 — failed to induce appreciable cell death in Ras-transformed MLBR cells at either concentration or were eliminated at this stage (**Figure 2**).

The deciding genotype was MLBR. Of the five lines in the panel, MLBR is the only one that combines BRCA1 depletion with oncogenic transformation, and it is therefore the closest available approximation to the disease state these compounds would need to act on: BRCA1-associated breast cancers are transformed tissue, not untransformed epithelium. A compound that kills BRCA1-depleted MLB cells but loses activity once the same cells acquire activated H-Ras is selective in the wrong direction — its window closes precisely where a therapeutic window would need to open. We therefore treated failure in MLBR as disqualifying regardless of performance elsewhere in the panel and applied the criterion at both concentrations so that a compound could not be retained on the strength of a saturating dose alone.

NSC98938, NSC300289, NSC275971, NSC367089 and NSC13051 showed heterogeneous effects, with one exception, unrelated to one another — NSC98938 is a curcuminoid, NSC300289 is the naphthalimide pinafide, NSC367089 is built on a naphthol AS-BI scaffold, NSC13051 is an acridine, and NSC275971 is a benzothiazol-6-yl carbamate. That structural heterogeneity is itself informative: the compounds that dropped out do not share a scaffold, which argues against their failure reflecting a single common liability and in favor of its reflecting the absence of the specific activity the screen was designed to detect. Curcuminoids are among the most frequently reported pan-assay interference scaffolds, showing activity across unrelated target classes through mechanisms including membrane perturbation, redox cycling and assay-signal interference rather than defined target engagement. Acridines and naphthalimides are classical DNA intercalators, a mode of action that produces broad cytotoxicity with little dependence on the repair status of the cell. Naphthol AS-BI is principally a chromogenic histochemical scaffold rather than a drug-like one.

### Secondary screening against diverse human cancer cells

To remove compounds whose activity reflected general cytotoxicity, the 13 hits were profiled by ATP-based CellTiter-Glo viability assay across a panel of human cancer lines at 0.1, 0.37, 1.1, 3.3 and 10 µM for 96 hours. Of the 13 *BRCA1*-knockdown–specific compounds entering secondary screening, eight were excluded in total by these criteria. The eliminated compounds are of interest but were not investigated in more detail in this report.

Five compounds survived both filters: NSC667251, NSC636948, NSC21683, NSC281383 and NSC136513 (**Table 2**, **Figure 3, 4**). All five retained activity in HCT116 *p53⁻/⁻* cells, providing independent confirmation of *p53*-independent killing.

**Figure 3.**
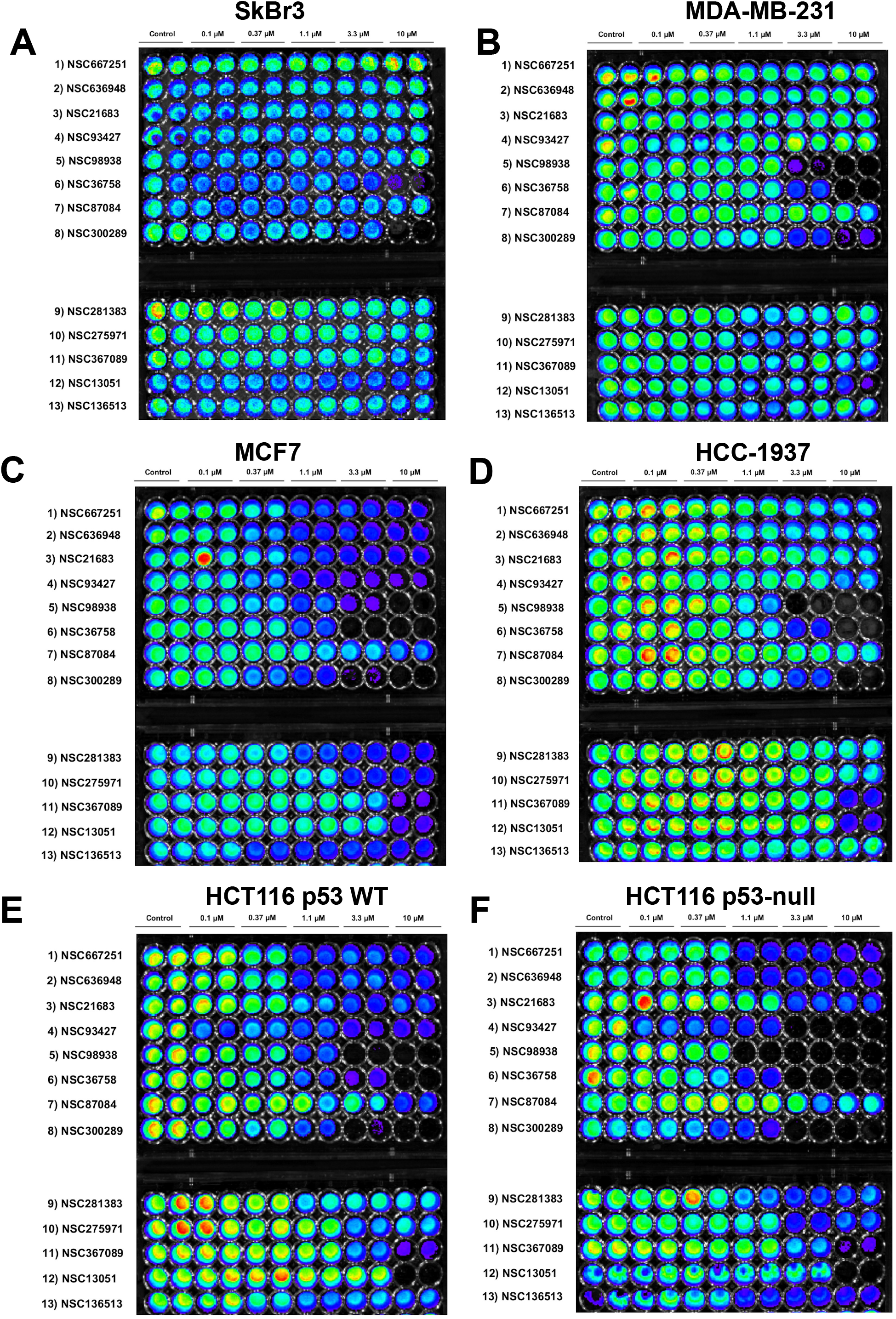
Dose-dependent growth inhibitory effects against human breast and colorectal cancer cell lines by 13 compounds chosen from primary screening based on suspected preferential activity towards BRCA1-knockdown isogenic cells. NCI DTP Diversity Set II NSC numbers are shown vertically in each panel, while the dose of each compound tested in duplicate is shown horizontally above each plate. Human cancer cell lines tested in this experiment were **(A)** SkBr3 breast cancer cells, **(B)** MDA-MB-231 breast cancer cells, **(C)** MCF7 breast cancer cells, **(D)** BRCA1-mutated HCC-1937 breast cancer cells, **(E)** HCT116 p53 WT colorectal cancer cells, and **(F)** HCT116 p53-null colorectal cancer cells.

**Figure 4.**
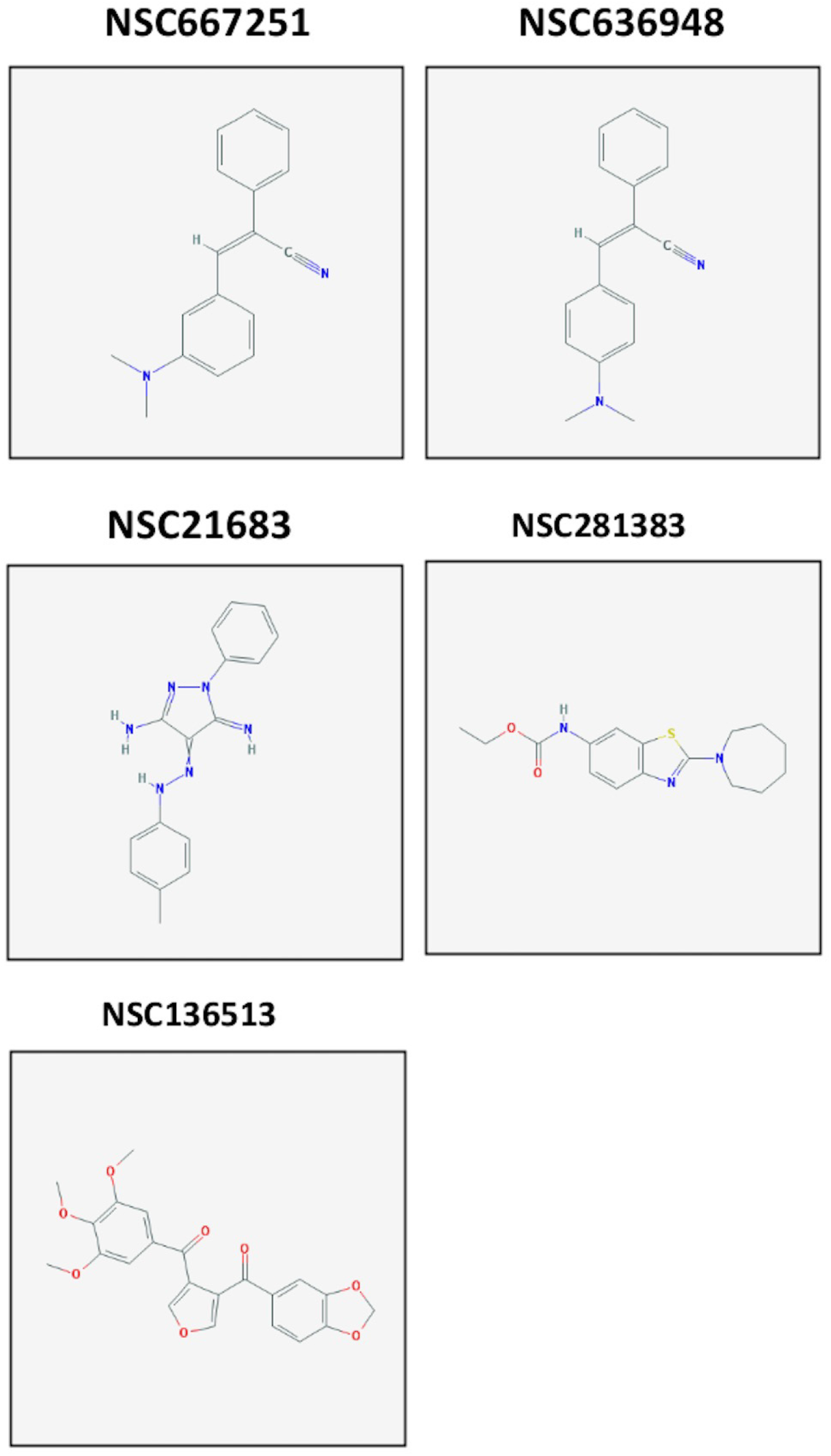
Molecular structures of five selected, BRCA1-knockdown specific small molecules after secondary screening against different cancer cells in Figures 2 and 3.

**Table 2.** Summary of small molecules selected for further characterization from the high throughput drug screen.

| Compounds Selected | Compounds Excluded |
| --- | --- |
| NSC667251<br>NSC636948<br>NSC21683<br>NSC281383<br>NSC136513 | NSC93427<br>NSC98938<br>NSC36758<br>NSC87084<br>NSC300289<br>NSC275971<br>NSC367089<br>NSC13051 |

### Sub-G1 analysis in normal, breast and ovarian cells demonstrates *BRCA1*-selectivity

The five selected compounds (NSC667251, NSC636948, NSC21683, NSC281383, and NSC136513) were evaluated by propidium iodide flow cytometry for sub-G1 DNA content — a direct measure of apoptotic nuclear fragmentation — across nine human cell lines treated at 5 µM for 96 hours (**Table 3**; **Figures 5-7**).

**Figure 5.**
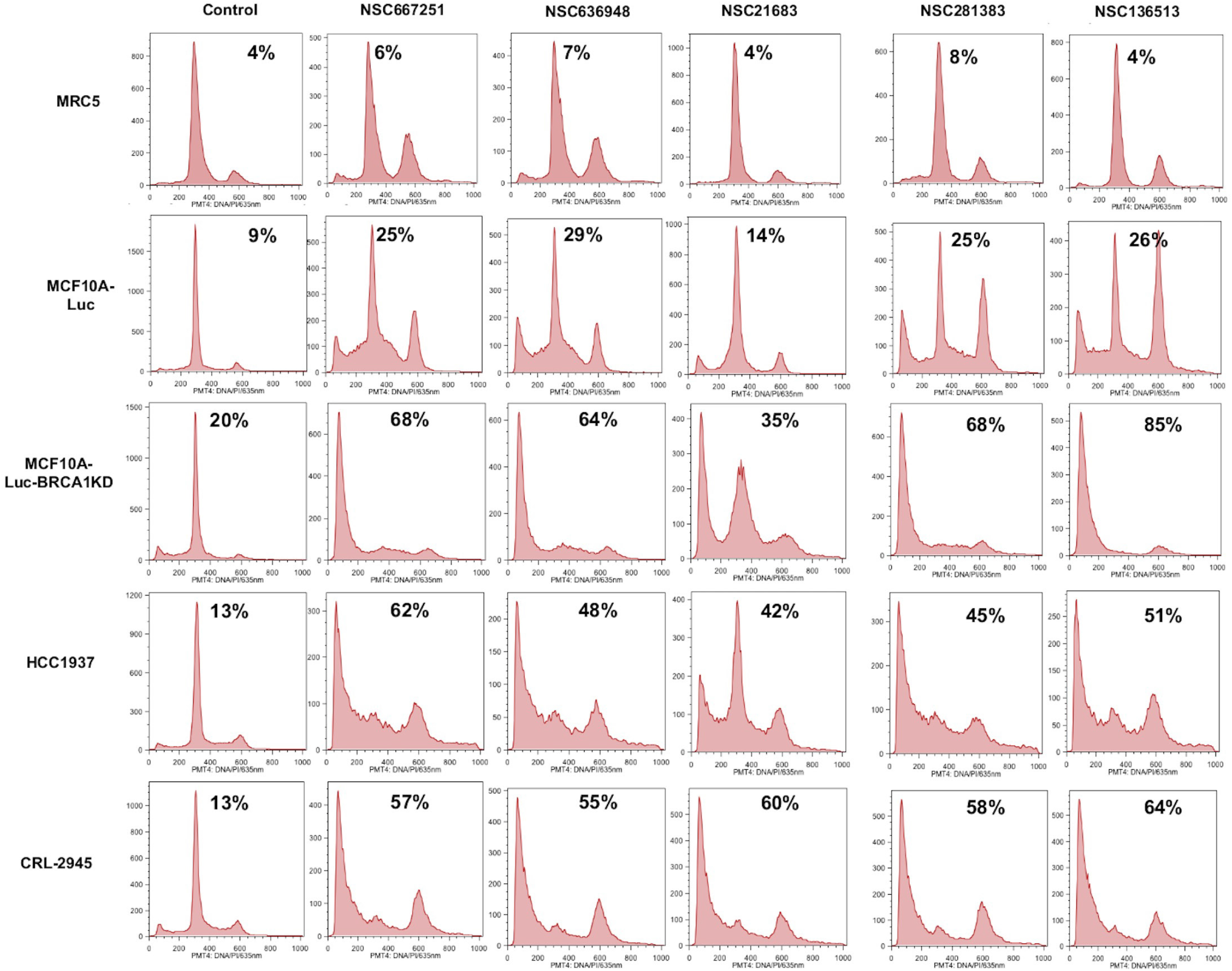
Demonstration of synthetic lethality by apoptosis induction in BRCA1-deficient human breast and ovarian cancer cells treated with isolated small molecule compounds. Each panel shows flow cytometric analysis of DNA content following propidium iodide staining. The sub-G1 content of the cell lines is listed numerically as a percent within each panel for each of the treatment conditions. Cell lines are listed to the left of the panels whereas the small molecules are listed above the panels along with the DMSO controls. MRC5 and MCF10A-Luc cells have wild-type BRCA1 and were tested as controls for comparison. Two human BRCA1-deficient breast cancer cell lines (MCF10A-Luc-BRCA1-KD and HCC-1937) and a human BRCA1-null ovarian cancer cell lines (CRL-2945) were tested as shown. HCC-1937 are homozygous for a pathogenic frameshift mutation in the BRCA1 gene (5382insC/c.5266dupC mutation) leading to two dysfunctional BRCA1 truncating allele. The MCF10A-Luc-BRCA1-KD cell line was derived from and is isogenic with MCF10A-luc except for BRCA1 knockdown. HCC-1937 and CRL-2945 are established cell lines derived from BRCA1-null human breast and ovarian tumors, respectively.

**Figure 6.**
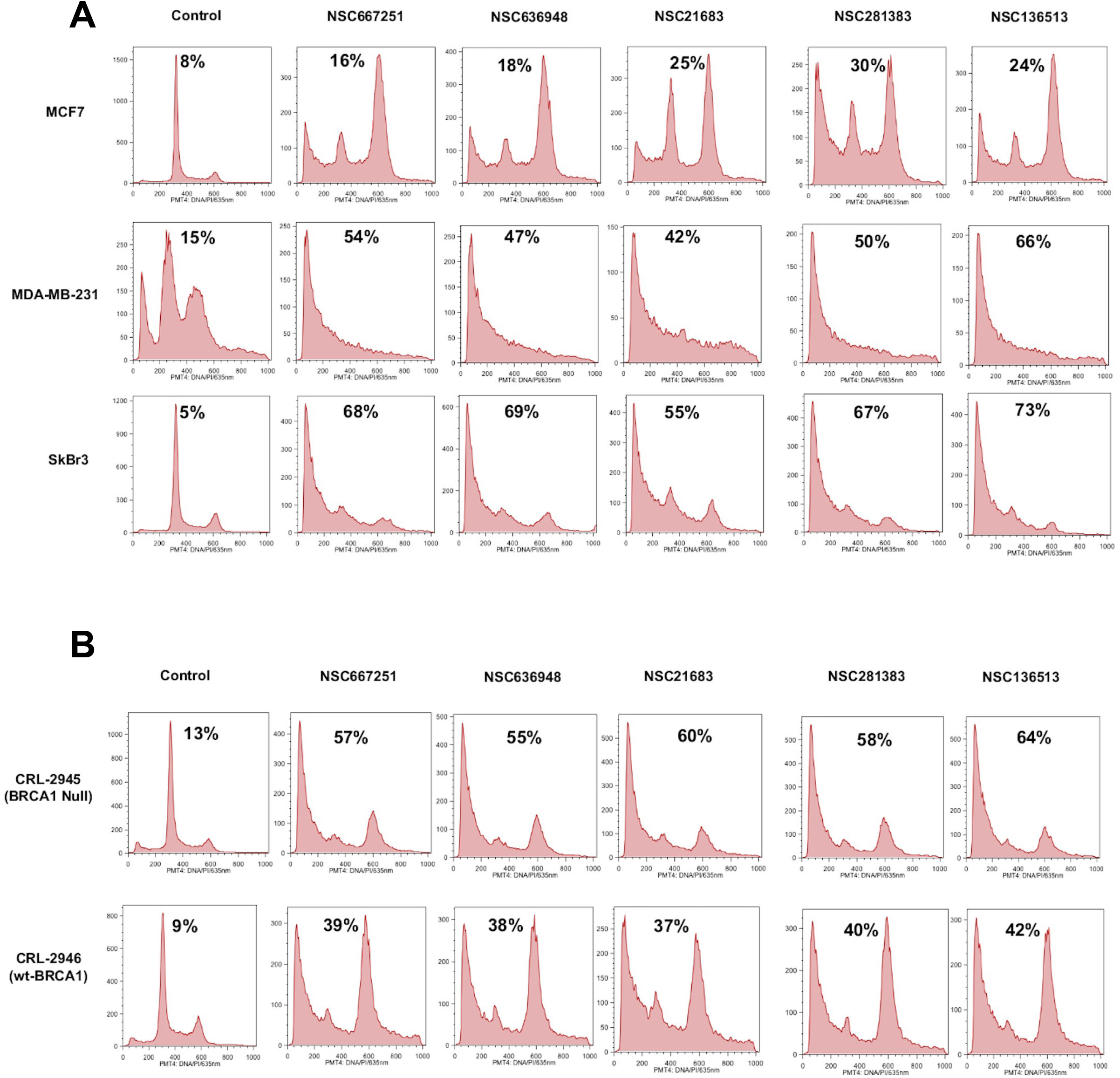
Apoptotic effects of small molecules against human breast and ovarian cancer cell lines. **(A)** Human breast cancer cells lines MCF7, MDA-MB-231 and SkBr3 were treated with small molecules isolated based on synthetic lethality phenotypes against BRCA1-deficient isogenic cell lines. **(B)** Human ovarian cancer cell lines CRL-2945 (BRCA1-null) and CRL-2946 (WT BRCA1) were treated with small molecules isolated based on synthetic lethality phenotypes against BRCA1-deficient isogenic cell lines. Each panel shows flow cytometric analysis of DNA content following propidium iodide staining. The sub-G1 content of the cell lines is listed numerically as a percent within each panel for each of the treatment conditions. Cell lines are listed to the left of the panels whereas the small molecules are listed above the panels along with the DMSO controls. Cells were treated with 5 μM of the compounds indicated for 96 hours before flow cytometric analysis.

**Figure 7.**
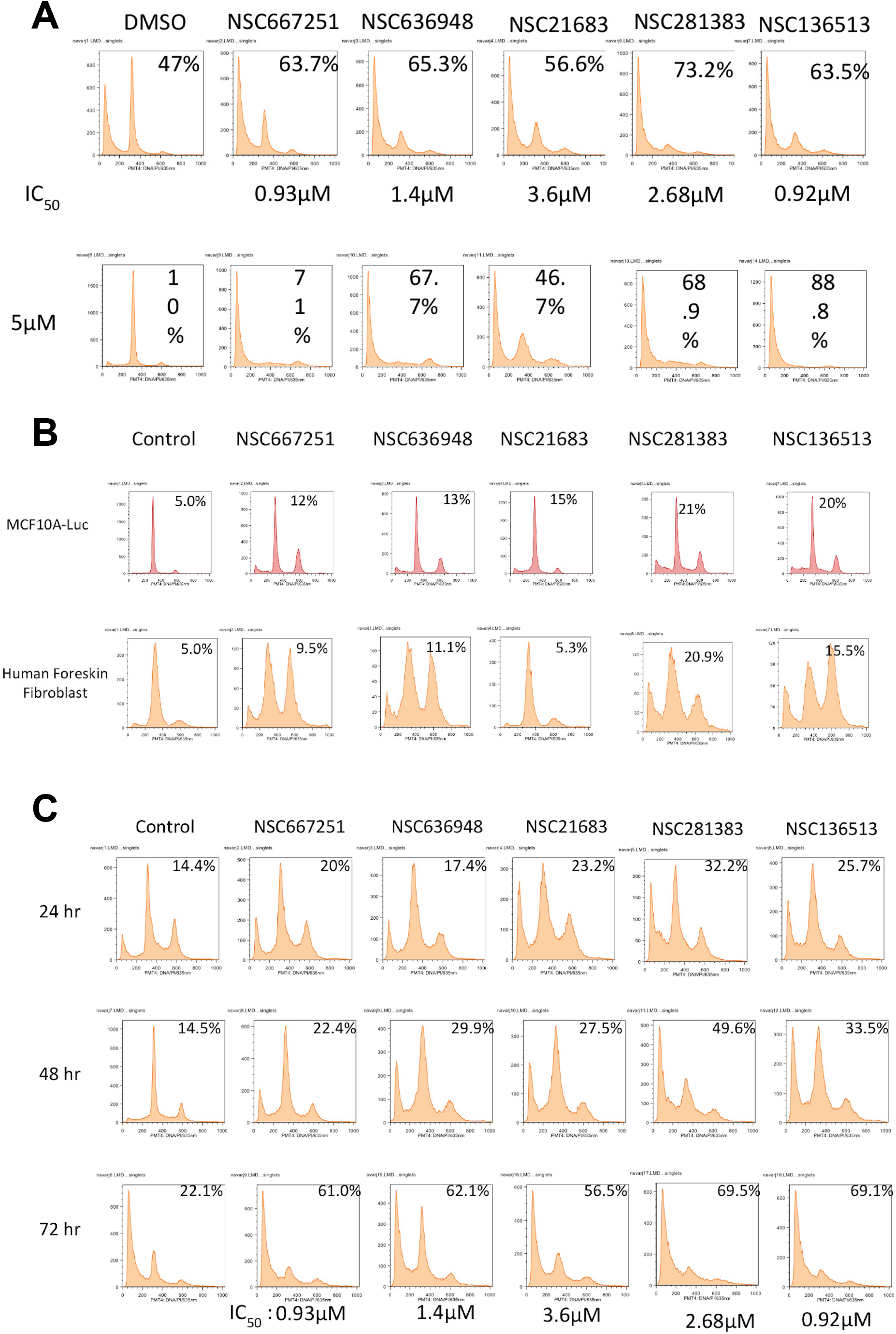
Proapoptotic effects of BRCA1-deficient synthetic lethal chemical compounds against cancer and normal cells. **(A)** Sub-G1 analysis of BRCA1-depleted MCF10A cells showing apoptotic cell death induction upon treatment for 96 hours with five small molecules individually at either IC50 dose as shown on top or 5 μM concentration as shown in the lower panels. **(B)** Sub-G1 analysis of normal human cells showing less apoptotic cell death induction upon treatment with five selected small molecules individually at 5 μM concentration for 96 hours. **(C)** Sub-G1 analysis of BRCA1-depleted MCF10A cells showing apoptotic cell death induction at different time intervals upon treatment with five small molecules individually at IC50 concentration for 24, 48, or 72 hours as indicated. The sub-G1 content of the cell lines is listed numerically as a percent within each panel for each of the treatment conditions. Cell lines are listed to the left of the panels whereas the small molecules are listed above the panels along with the DMSO controls.

**Table 3.** Summary of different, human normal and cancer cells tested for the anti-tumorigenic potential of five selected small molecules at 5 µM concentration for 96 hrs.

| No | Cell Lines | % Cells in Sub G1 |  |  |  |  |  |
| --- | --- | --- | --- | --- | --- | --- | --- |
|  |  | Control | NSC667251 | NSC636948 | NSC21683 | NSC281383 | NSC136513 |
| 1 | MRC5 | 4 | 6 | 7 | 4 | 8 | 4 |
| 2 | MCF10A-Luc (ML) | 9 | 25 | 29 | 14 | 25 | 26 |
| 3 | ML-BRCA1KD (MLB) | 20 | 68 | 64 | 35 | 68 | 85 |
| 4 | HCC1937 | 13 | 62 | 48 | 42 | 45 | 51 |
| 5 | MDA-MB-231 | 15 | 54 | 47 | 42 | 50 | 66 |
| 6 | SkBr3 | 5 | 68 | 69 | 55 | 67 | 73 |
| 7 | MCF7 | 8 | 16 | 18 | 25 | 30 | 24 |
| 8 | OC-2945(Null) | 13 | 57 | 55 | 60 | 58 | 64 |
| 9 | OC-2946(+BRCA1) | 9 | 39 | 38 | 37 | 40 | 42 |

**Table 4.**
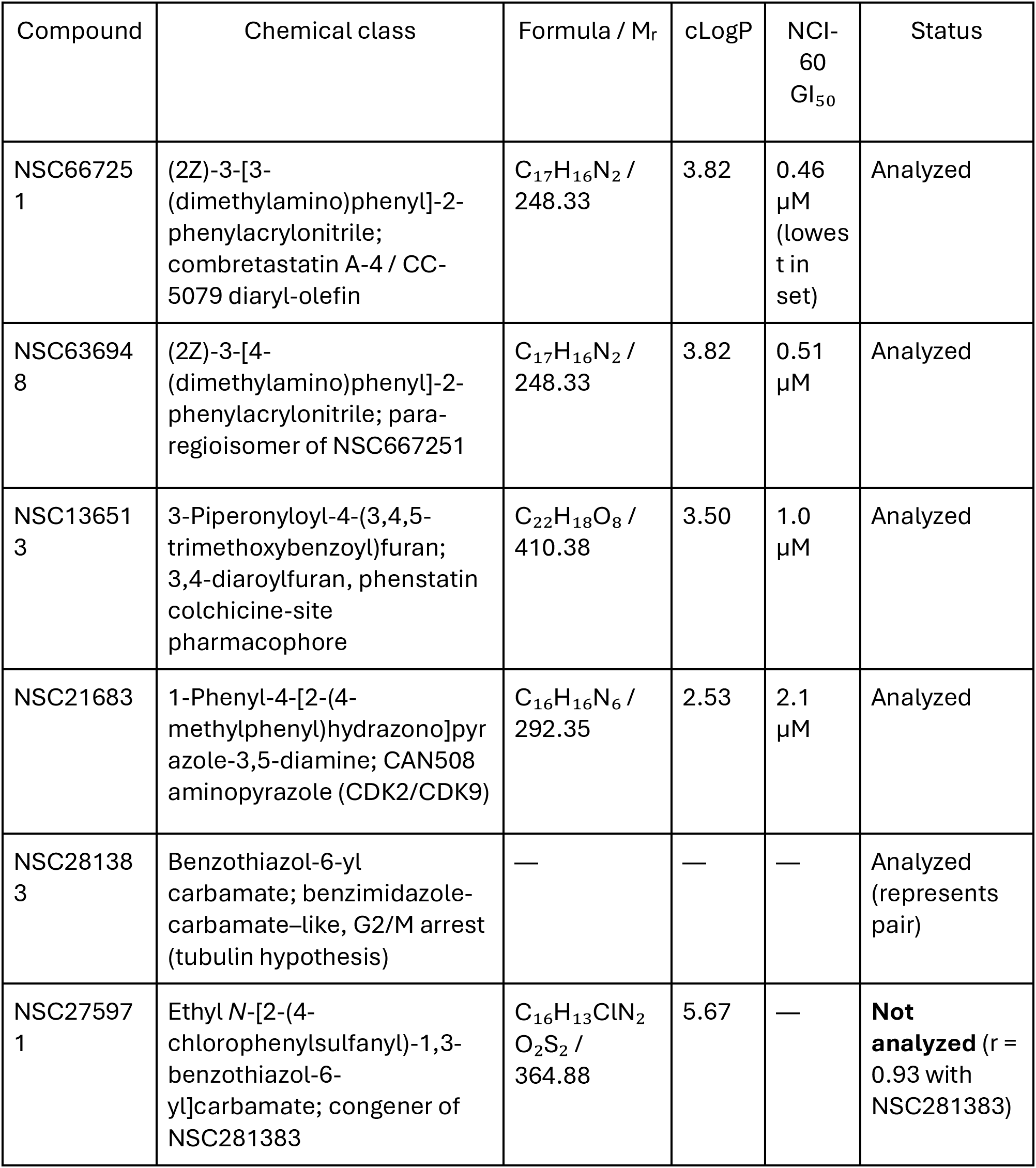
Chemotype, physicochemical properties and NCI-60 potency of the six CellMiner family members.

Three patterns of interest emerged from this dataset. First, the cytotoxicity response tracks *BRCA1*-status. In the isogenic mammary pair, apoptosis rose from 9% in ML cells to 20% baseline and 35–85% after treatment in MLB cells, and BRCA1-mutant HCC1937 cells responded comparably (42–62%). The most direct test is the isogenic ovarian pair: *BRCA1*-null CRL-2945 cells reached 55–64% sub-G1, whereas the same cells reconstituted with wild-type BRCA1 (CRL-2946) reached only 37–42% — a consistent reduction of roughly one-third for every compound, achieved by restoring a single gene (**Figure 5**). Second, normal cells are largely spared. MRC5 normal fibroblasts remained at 4–8% sub-G1, essentially indistinguishable from the 4% vehicle control, and this was confirmed in a separate panel of normal human foreskin fibroblasts and MCF10A-Luc cells (**Figure 5**), where responses ranged from 5.3% to 20.9% against a 5.0% control. MCF7 cells, which are wild-type for both *BRCA1* and *TP53*, were similarly resistant (16– 30%). Third, killing does not require p53. TP53-mutant MDA-MB-231 (42–66%) and SKBr3 (55– 73%) cells were among the most sensitive lines tested, exceeding the response of *TP53*–wild-type MCF7 cells by two-to four-fold (**Figure 6A**). Taken with the HCT116 p53⁻/⁻ viability data and the MLBP screening data, this establishes that apoptotic induction proceeds through a p53-independent route.

### Potency and kinetics of apoptotic induction by novel *BRCA1*-selective therapeutic agents

Half-maximal inhibitory concentrations determined in BRCA1-depleted MCF10A cells were 0.93 µM (NSC667251), 1.4 µM (NSC636948), 3.6 µM (NSC21683), 2.68 µM (NSC281383) and 0.92 µM (NSC136513). When cells were treated at each compound’s own IC₅₀ rather than at a uniform 5 µM, comparable levels of apoptosis were achieved (**Figure 7**), indicating that the differences observed in **Table 3** largely reflect differences in potency rather than in maximal efficacy.

Time-course analysis at IC₅₀ doses revealed two distinct kinetic classes (**Figure 7**). NSC281383 and NSC136513 acted rapidly, reaching 32.2% and 25.7% sub-G1 by 24 hours and 49.6% and 33.5% by 48 hours, against a 14.4–14.5% vehicle background. NSC667251, NSC636948 and NSC21683 were slower, remaining at 22.4–29.9% at 48 hours. By 72 hours the two classes had converged, with all five compounds producing 56.5–69.5% sub-G1 versus 22.1% in controls. All five therefore reach a similar apoptotic endpoint, but by different routes — a divergence that both informed the design of the immunoblot experiments below and may prove useful in designing combination regimens.

### γ-H2AX accumulation identifies *BRCA1*-selective therapeutic compounds as genotoxic agents

To determine whether apoptosis was preceded by DNA damage, we assessed phosphorylation of histone H2AX. H2AX is a variant of core histone H2A that constitutes a minor fraction of the nucleosomal H2A pool but is distributed throughout chromatin (**Figure 8, 9**). Within minutes of a DNA double-strand break, the phosphatidylinositol 3-kinase–related kinases ATM, ATR and DNA-PK proteins phosphorylate H2AX on serine 139 of its C-terminal SQEY motif, generating the modified form designated γ-H2AX. This modification is not confined to the break itself: it propagates across megabase-scale domains of chromatin flanking the lesion, amplifying a single molecular event into a structure large enough to be resolved by light microscopy. Each discrete nuclear focus therefore corresponds to one double-strand break, and focus counts provide a quantitative index of the physical integrity of nuclear DNA. H2AX foci serve as an assembly platform recruiting MDC1, the MRE11–RAD50–NBS1 complex, 53BP1 and BRCA1 to catalyze DNA repair. In cells with intact homologous recombination, H2AX foci resolve as breaks are repaired, so persistence of γ-H2AX signal beyond the expected repair window indicates lesions that the cell cannot resolve or are not repaired. In *BRCA1*-deficient cells, where homologous recombination is compromised, this is precisely the expected outcome and is what was observed in our studies.

**Figure 8.**
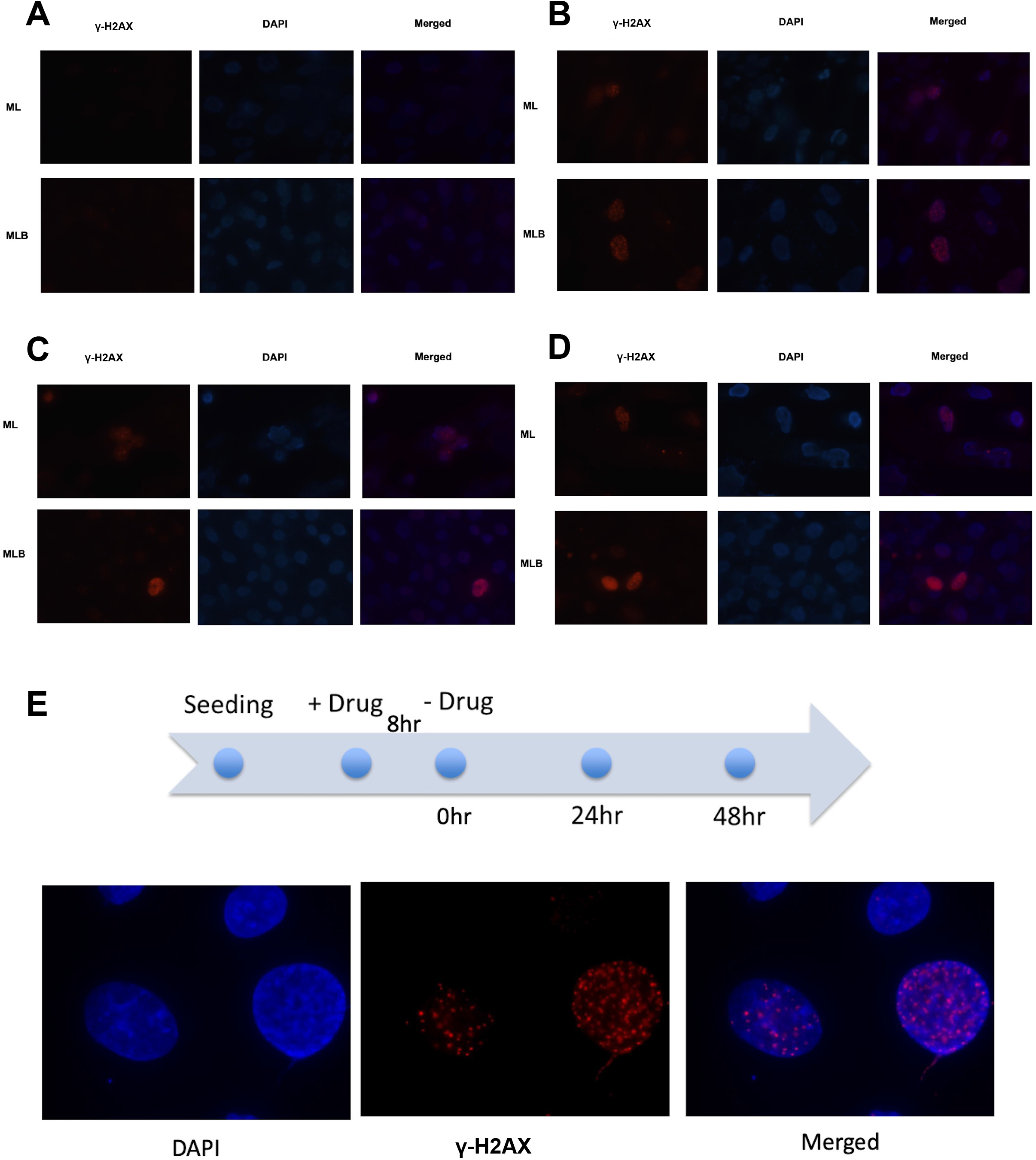
Assessment of drug-induced DNA damage in control MCF10A-Luc (ML), BRCA1-deficient MCF10A-Luc-BRCA1-KD (MLB) of BRCA1-deficient human HCC-1937 breast cancer cells. (A) DNA damage analysis of BRCA1-depleted MCF10A cells (MLB) versus control isogenic ML cells treated with DMSO followed by γ-H2AX staining showing no genotoxicity. (B) DNA damage analysis of BRCA1-depleted MCF10A cells (MLB) versus control isogenic ML cells treated with small molecule drug NSC21683 at 5 μM for 20 hours followed by γ-H2AX staining that shows greater immunofluorescent staining indicative of DNA double-strand break damage induction in MLB versus ML cells. (C) DNA damage analysis of BRCA1-depleted MCF10A cells (MLB) versus control isogenic ML cells treated with small molecule drug NSC281383 at 5 μM for 20 hours followed by γ-H2AX staining that shows greater immunofluorescent staining indicative of DNA double-strand break damage induction in MLB versus ML cells. (D) DNA damage analysis of BRCA1-depleted MCF10A cells (MLB) versus control isogenic ML cells treated with small molecule drug NSC136513 at 5 μM for 20 hours followed by γ-H2AX staining that shows greater immunofluorescent staining indicative of DNA double-strand break damage induction in MLB versus ML cells. (E) DNA damage analysis of HCC-1937 cells treated with small molecule drug NSC21683 at 5 μM for 8 hours and the cells were then changed to drug free media. γ-H2AX staining shows the persistence of DNA damage even after 72 hours of releasing the BRCA1-deficient HCC-1937 dells to drug free media.

**Figure 9.**
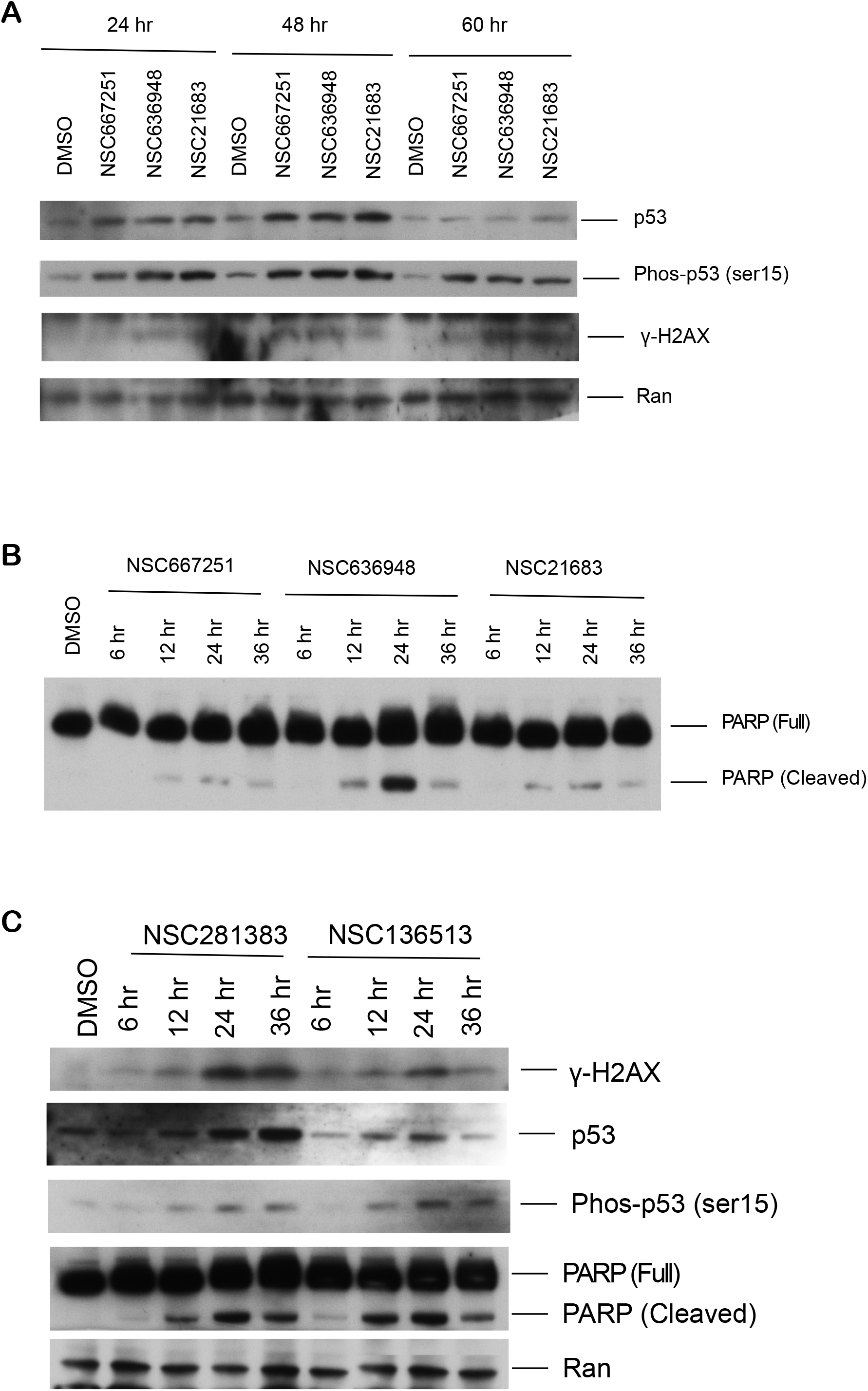
Induction of a DNA damage response markers and cell death on BRCA1-depleted MCF10A cells (MLB) by small molecules chosen for their synthetic lethal phenotype in BRCA1-deficient cells. (A) Western blot analysis of BRCA1-depleted MCF10A cells treated with NSC667251, NSC636948, and NSC21683 at a 5 μM dose for different time intervals as indicated. The results show increased levels of p53, phosphorylated-p53, and γ-H2AX upon drug treatment indicative of induction and persistence of DNA damage signaling. Ran was used as a protein loading control. **(B)** Western blot analysis of BRCA1-depleted MCF10A cells treated with NSC667251, NSC636948, and NSC21683 at a 5 μM dose for different time intervals as indicated. The results show a time dependence to PARP cleavage consistent with apoptosis induction. Full-length PARP is used as an internal loading control for this blot.

Immunofluorescence staining of ML and MLB cells treated for 20 hours at 5 µM showed minimal γ-H2AX signal under vehicle conditions in either genotype (**Figure 8**). Treatment with NSC21683 (**Figure 8B**), NSC281383 (**Figure 8C**) or NSC136513 (**Figure 8D**) produced abundant, discrete γ-H2AX nuclear foci that colocalized with DAPI-stained chromatin, and in every case the foci were substantially more numerous and more intense in BRCA1-depleted MLB cells than in isogenic ML controls. The persistence of these lesions was tested directly in BRCA1-mutant HCC1937 cells: after only 8 hours of exposure to NSC21683 at 5 µM followed by transfer to drug-free medium, γ-H2AX foci remained clearly detectable 72 hours after washout (**Figure 8E**), demonstrating that the damage inflicted is not efficiently reversed once the compound is removed.

### NSC667251, NSC636948, NSC21683, NSC281383, and NSC136513 induce DNA damage signaling in *BRCA1*-deficient cells

NSC281383 and NSC136513 were characterized by immunofluorescence instead of western analysis (**Figures 8C and 8D**). This was felt to be the appropriate assay for their kinetics allowing use of immunofluorescence as a single-cell readout in which damaged and undamaged nuclei are scored individually. Thus, the presence of an apoptotic subpopulation would not be expected to confound the measurement, and the short 20-hour exposure used for staining the cells falls inside the window before either compound produces substantial cell death. NSC21683 was deliberately included in both formats and serves as a bridging control, confirming that the two readouts report concordantly on the same compound.

Western blot analysis of MLB lysates confirmed the immunofluorescence findings biochemically. Treatment with NSC667251, NSC636948, NSC21683, NSC281383, or NSC136513 at 5 µM induced γ-H2AX that was detectable at 24 hours and persisted through 48 and 60 hours, alongside stabilization of total p53 and accumulation of phospho-p53 (Ser 15), with Ran as loading control (**Figure 9A, C**). The five compounds produced cleavage of poly(ADP-ribose) polymerase as early as 12 hours after treatment (**Figure 9B, C**). Across the immunofluorescence and immunoblot datasets taken together, γ-H2AX induction was documented for all five compounds, establishing DNA damage as a shared property of the series. That p53 was stabilized and phosphorylated yet dispensable for killing (**Table 3**; **Figure 9A, C**) indicates that the damage response is engaged but that execution proceeds through p53-independent effectors.

### NSC636948 is a PPARβ/δ-directed ligand, and *PPARD* expression is prognostic in breast cancer

Among the five compounds carried forward, NSC636948 is the only member of the series with a directly annotated molecular target, and it was therefore selected for focused mechanistic follow-up. NSC636948 and NSC667251 emerged together from the primary screen as positional isomers of a single scaffold: both are built on a phenylacrylonitrile core bearing a dimethylamino-substituted phenyl ring, and they differ only in the position at which the dimethylamino group attaches. NSC636948 is the para form, [(Z)-3-(4-(dimethylamino)phenyl)-2-phenylacrylonitrile] (C₁₇H₁₆N₂, Mᵣ 248.33); NSC667251 is the meta regioisomer.

To assess if the identified compounds also share similar toxicity in an orthogonal, database-derived dataset, NCI-60 growth-inhibition profiles for all 13 *BRCA1*-selective hits from CellMiner were retrieved. Pairwise correlations were then computed based on the activity fingerprints of the compounds. Six compounds formed a coherent family with mean within-group correlations of r= 0.53-0.72; no other compounds exceeded r= 0.5 by the same measure, affirming their classification as distinct in their responses across cancer types, as observed in *in vitro* cytotoxicity analyses (**Figure 10**). Aggregated across the nine NCI-60 tissue panels, this family showed a consistent and specific tissue signature: with a pronounced sensitivity to leukemia, colon cancer, and melanoma, and a relative resistance to renal, ovarian and lung cancer.

**Figure 10.**
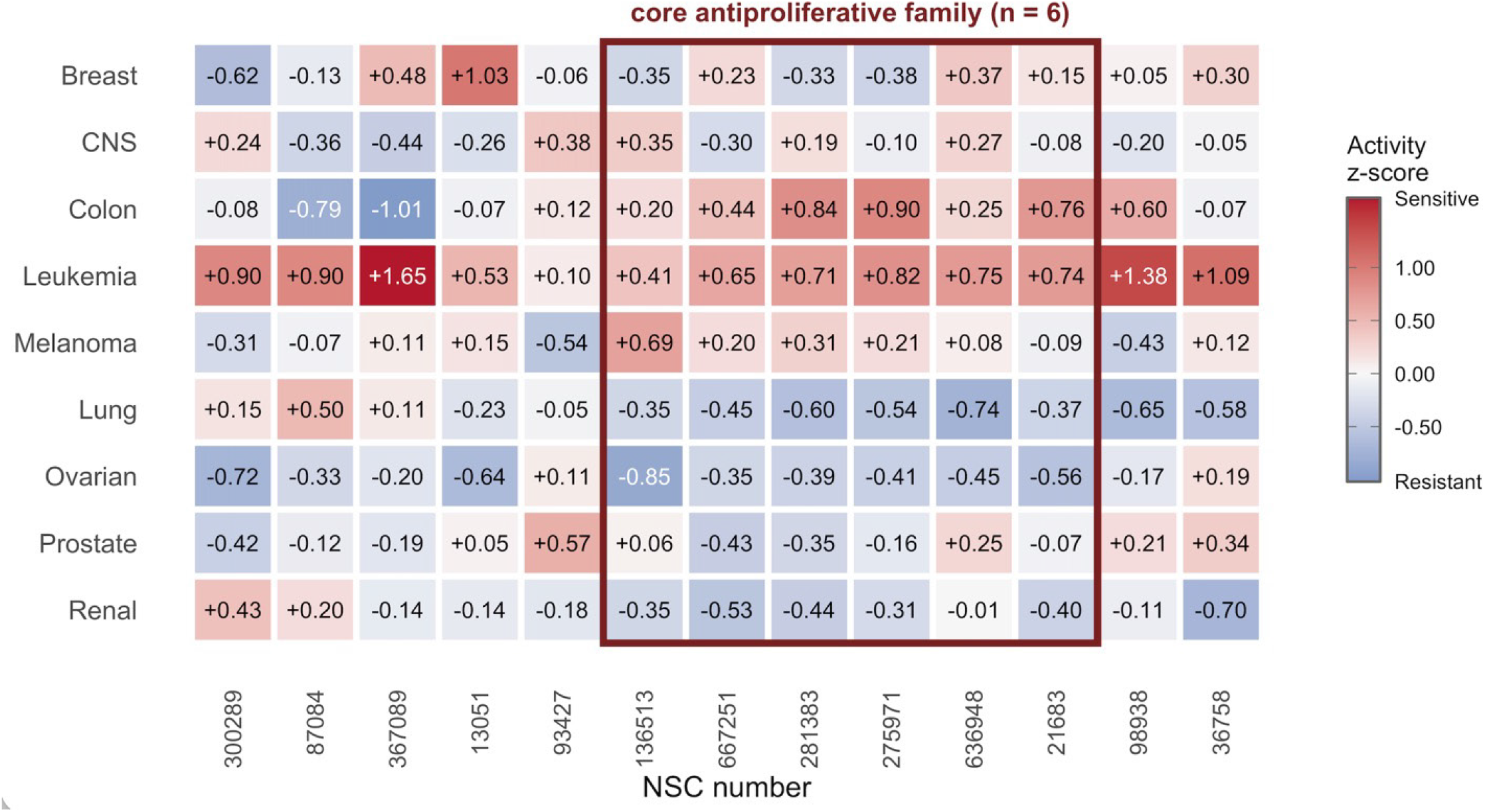
NCI-60 tumor tissue-of-origin activity profile for 13 screened *BRCA1*-selective compounds. Heatmap of mean compound activity z-scores across nine NCI-60 tissue-of-origin subpanels (breast, n = 5; CNS, n = 6; colon, n = 7; leukemia, n = 6; melanoma, n = 10; lung, n = 9; ovarian, n = 7; prostate, n = 2; renal, n = 8 cell lines) for 13 compounds (NSC300289, NSC87084, NSC367089, NSC13051, NSC93427, NSC136513, NSC667251, NSC281383, NSC275971, NSC636948, NSC21683, NSC98938, NSC36758). Per-cell-line activity z-scores were derived from GI50 values (sulforhodamine B assay [45]) obtained from the NCI Developmental Therapeutics Program via CellMiner v2.15 [46] and averaged within each tissue group; cell lines lacking a validated value for a given compound were excluded from that average. Red indicates relative sensitivity (positive z-score); blue indicates relative resistance (negative z-score). Columns outlined in red (NSC136513, NSC667251, NSC281383, NSC275971, NSC636948, NSC21683) denote the six compounds comprising the “core antiproliferative group” (n = 6). These six compounds were defined by pairwise Pearson correlation of each compound’s NCI-60 activity z-score pattern against the other members of the group. All six showed mean within-group correlations of r = 0.53–0.72; no other screened compound exceeded r = 0.5 by the same measure (maximum r = 0.43, NSC93427).

Independent work has established that NSC636948 is broadly cytotoxic. Among a series of (Z)-2,3-diphenylacrylonitrile analogues it showed the highest activity across lung (A549), ovarian (SKOV-3), melanoma (SK-MEL-2) and colon (HCT15) models [35]. This is not in conflict with the tissue signature reported in **Figure 10**: NCI-60 mean-graph z-scores are normalized within each compound’s own activity profile and therefore describe relative rather than absolute tissue preference, so a compound may be potent across all panels while still ranking a given tissue below its own mean.

The molecular target of NSC636948 was subsequently identified as PPARβ/δ, establishing this compound as a selective inverse agonist of the receptor [36]. NSC636948, together with its positional isomer NSC667251, selectively binds PPARβ/δ with minimal cross-reactivity toward PPARα and PPARγ and represses PPARβ/δ-dependent transcription. These studies established the diarylacrylonitrile scaffold as a platform for selective PPARβ/δ inhibition, leading to the development of related compounds, including DG172 [36]. The identification of active PPARβ/δ ligands across this scaffold class demonstrated that pharmacological repression of PPARβ/δ signaling is feasible and may provide a therapeutic strategy for targeting tumor-promoting metabolic and transcriptional programs.

PPARs are ligand-activated nuclear receptors that regulate transcription in response to fatty acids and eicosanoid metabolites, binding as heterodimers with retinoid X receptor to PPAR response elements and acting through both transcriptional activation and repression [41,42]. Of the three isoforms, PPARα governs lipid metabolism in liver and skeletal muscle and PPARγ controls adipocyte differentiation, lipid storage and glucose homeostasis — both are established drug targets, for dyslipidemia and diabetes respectively — whereas PPARβ/δ carries broader responsibilities spanning fatty acid oxidation, proliferation, differentiation, apoptosis, angiogenesis, inflammation and tumorigenesis [37–40,43]. Its wide tissue distribution and its capacity to redirect lipid utilization under cellular stress make it an integrator of metabolic and inflammatory signaling, and a growing body of evidence identifies it as an attractive therapeutic target in both metabolic disease and cancer.

This is directly relevant to the *BRCA1*-deficient setting. Elevated fatty acid oxidation contributes to progression in cancers of the liver, lung, breast and prostate, and as a lipid-sensing receptor. PPARβ/δ controls fatty acid uptake, oxidation and stress-response programs that allow tumor cells to survive nutrient deprivation and hypoxia. Homologous recombination–deficient tumors are initially sensitive to PARP inhibition, but PPARβ/δ-driven fatty acid oxidation regenerates intracellular NAD⁺ and can thereby sustain the DNA-repair capacity on which resistance depends. Antagonism of PPARβ/δ would be expected to close that route, which makes the recovery of a PPARβ/δ inverse agonist from a BRCA1-selectivity screen mechanistically coherent rather than incidental.

Clinically, elevated expression of PPARβ/δ and its target genes has been associated with unfavorable outcomes across multiple tumor types, and in breast cancer specifically (**Figure 11A**), increased PPARD expression and activity have been linked to more aggressive disease phenotypes including triple-negative and HER2-positive subtypes [41,44]. Our own analysis reproduces this subtype distribution, showing that *PPARD* gene expression, which encodes the nuclear receptor PPARβ/δ, was highest in basal-like (median ∼2.93) and HER2-enriched (median ∼2.91) tumors, and lower in luminal A and luminal B disease (median ∼2.87 for both) (**Figure 11C**). The enrichment in the basal-like compartment is notable (**Figure 11D**), since that is the transcriptional subtype in which *BRCA1*-mutant tumors overwhelmingly arise.

**Figure 11.**
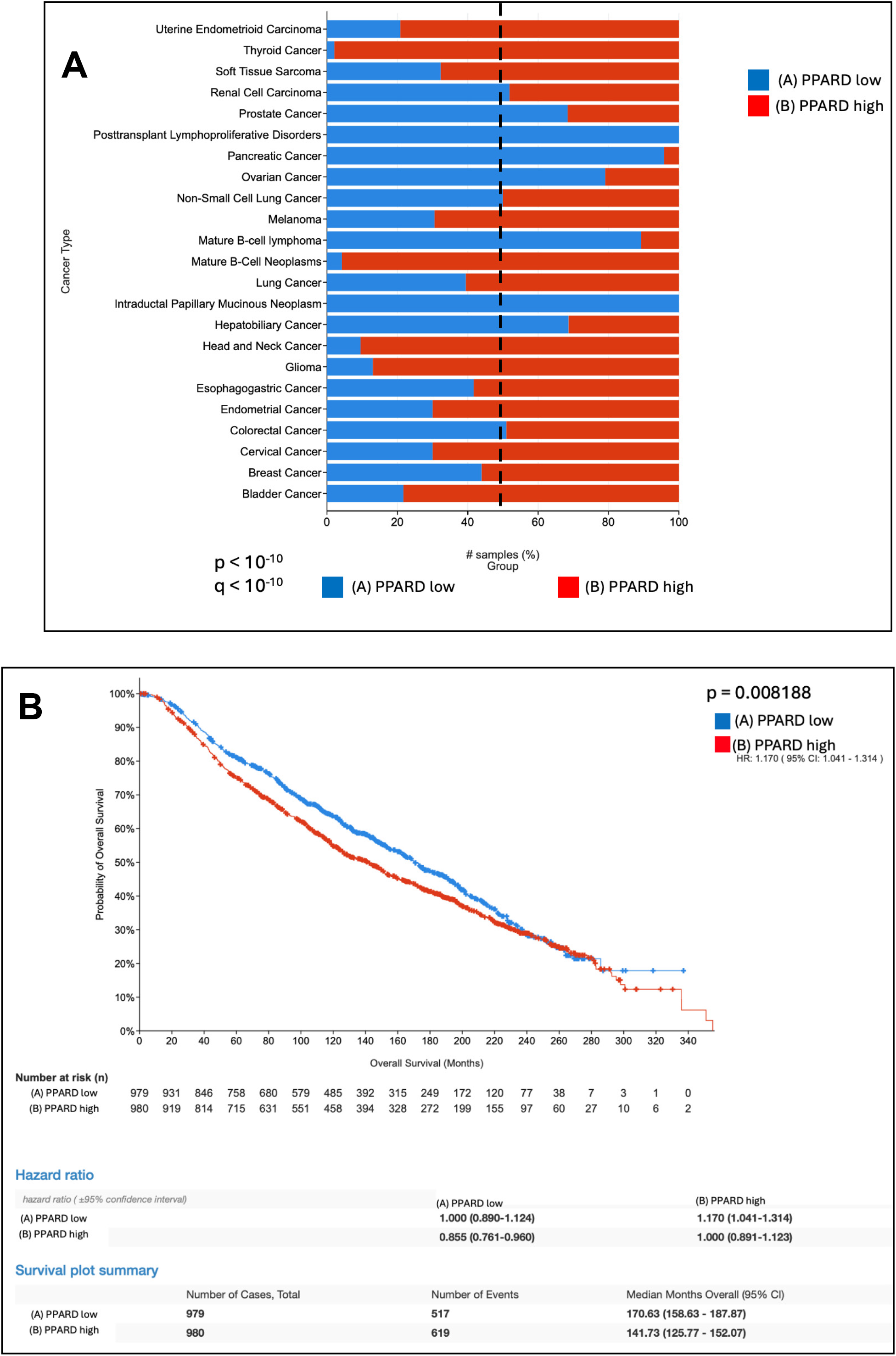
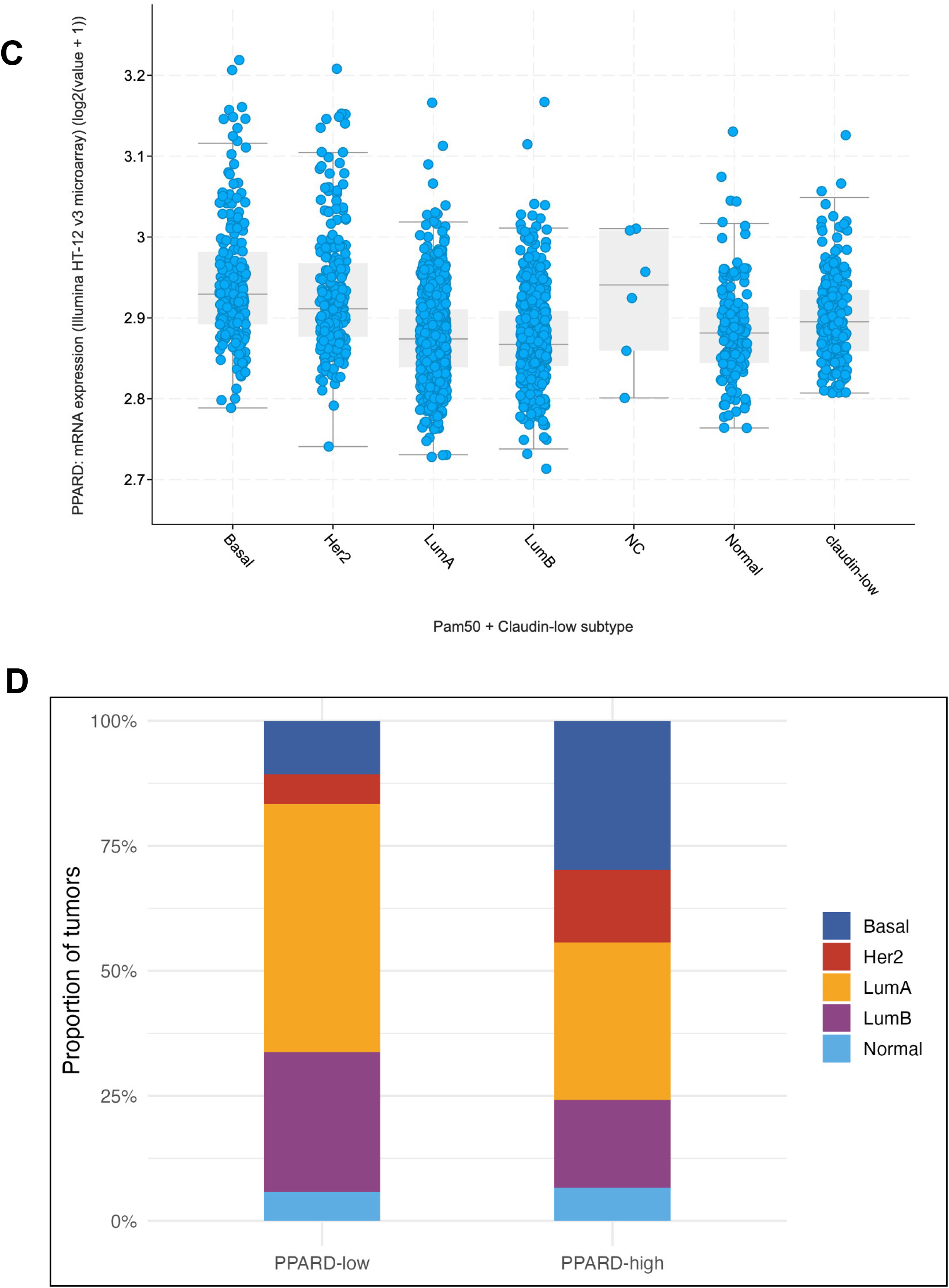
PPARD expression in cancer including breast cancer subtypes and impact on survival. (A) PPARD expression in TCGA pan-cancer atlas. Patterns of *PPARD* expression across cancer types. (B) Elevated PPARD mRNA expression in breast cancer correlates with poor patient survival. K-M survival plot between the lower median (blue, A) and higher median (red, B) *PPARD* mRNA expression in the breast cancer cohort. (C). *PPARD* mRNA expression levels across breast cancer molecular subtypes. Data is derived from Illumina Human HT-12 v3 microarray platform. Tumor subtypes analyzed: Basal – basal-like; HER2 – HER2 enriched; LumA – luminal A; LumB – luminal B; NC – non-cancerous control; Normal – normal-like; claudin-low – claudin-low breast cancer samples. (D) Distribution of PAM50 molecular subtypes within *PPARD*-low and *PPARD*-high breast tumors. This 100% stacked bar chart illustrates the proportional distribution of PAM50 intrinsic molecular subtypes within PPARD-low and PPARD-high breast tumors groups. Data is aggregated from a combined dataset (METABRIC + TCGA datasets, total 2955 patients).

Consistent with these observations, Kaplan–Meier analysis of a combined breast cancer cohort of 1,959 patients assembled from the METABRIC datasets demonstrated that patients with high PPARD expression, stratified at the median, had significantly worse OS than those with low expression (HR 1.170, 95% CI 1.041–1.314; log-rank p = 0.008) (**Figure 11B**). Elevated PPARD therefore behaves as a negative prognostic biomarker in breast cancer, and its concentration within the basal-like subtype places the target in the same patient population as the BRCA1 defect that this compound series was selected to exploit.

### NSC21683 is a less potent *BRCA1*-selective agent

The IC₅₀ of NSC21683 of 3.6 µM in *BRCA1*-depleted cells is 2.4-fold above the mean of the other four compounds (1.48 µM) and 3.9-fold above that of the most potent member, NSC136513 (0.92 µM), and this ranking is reproduced independently in the NCI-60 data, where its GI₅₀ of 2.1 µM is the highest in the family (**Table 5**). The same pattern runs through every cellular assay: NSC21683 produced the lowest sub-G1 response of the series in *BRCA1*-depleted MLB cells (35% versus 64–85% for the others; **Table 3**), the lowest response in SKBr3 cells (55%), and the poorest discrimination between *BRCA1*-deficient and *BRCA1*–wild-type breast cells, with an MLB:MCF7 ratio of 1.4 compared with 2.3–4.3 for the remainder. Its position at the edge of the chemical group is consistent with this. NSC21683 is the only member whose annotated target — CDK2/CDK9, via the CAN508 aminopyrazole chemical type — lies outside the tubulin-directed axis shared by the other chemicals. NSC21683 nonetheless retains gross and not unexpected mechanistic features of the series, showing robust γ-H2AX focus formation with damage persisting 72 hours after washout (**Figures 5B and 5E**) and PARP cleavage by 12 hours (**Figure 5G**), and its activity against *BRCA1*-null CRL-2945 ovarian cells (60%) is comparable to that of the other compounds. It is therefore best regarded as the least potent member of the identified set, and as the compound least likely to be advanced without medicinal-chemistry optimization.

### NSC281383 is a more toxic and less *BRCA1*-selective agent

NSC281383 produced the highest apoptotic response of the series in every normal or repair-proficient cell type examined: 8% in MRC5 fibroblasts (versus 4% control and 4–7% for the other compounds), 30% in *BRCA1/TP53*–wild-type MCF7 cells (versus 16–25%), and 20.9% in normal human foreskin fibroblasts and 21% in normal MCF10A-Luc cells (versus 5.0% control; **Figure 7B**) — roughly four-fold over background, and the highest normal-cell toxicity recorded in that panel. Consequently, although its potency against *BRCA1*-deficient cells is good (IC₅₀ 2.68 µM; 68% sub-G1 in MLB) and its apoptotic onset is the fastest of the series (32.2% sub-G1 by 24 hours; Figure 7C), its selectivity window is the narrowest: the MLB-to-normal-fibroblast ratio is 3.3, compared with 5.5–7.2 for the other four compounds. Rapid onset combined with the least favorable therapeutic index suggests a mechanism that is not fully contingent on *BRCA1* loss. This concern extends to its structural class, since the benzothiazole carbamate scaffold is subject to carbamate hydrolysis to an arylamine, and the closely related NSC275971 releases 2-mercaptobenzothiazole; any further development of NSC281383 should therefore be preceded by a formal assessment of both its selectivity margin and its metabolic liabilities.

## Discussion

*BRCA1* dysfunction is associated with the existence of a distinctive subgroup of breast cancer cases exhibiting severe impairments in the DNA repair process and genomic instability [10–13]. These defects make the affected tumors susceptible to various forms of chemotherapy, especially PARP inhibition and platinum drugs; however, drug resistance is a formidable challenge when attempting long-term cancer control [20–27]. In this study, we report five compounds that cause BRCA1-deficient breast tumor cells to undergo apoptosis.

A key strength of our study is use of an isogenic human mammary epithelial cell system, which is genetically defined, by contrast to previously utilized tumor cells that have undergone significant genetic modifications. Although activating *KRAS* or *HRAS* mutations are uncommon in *BRCA1*-associated TNBC, H-RasG12V expression was used as a transformation strategy to generate a tumorigenic isogenic model while preserving the *BRCA1*-deficient genetic background. The outcome of this method was the identification of drugs that kill cells via the loss of *BRCA1* function.

The compounds identified in this screen consistently induced apoptosis in *BRCA1*-mutant cells through activation of DNA damage responses, which was confirmed by the presence of phosphorylated histone γ-H2AX as well as PARP cleavage early. Importantly, apoptosis was observed in TP53-mutant and TP53-null cellular models, suggesting that functional p53 is not essential for compound activity. This observation is clinically relevant because *TP53* is frequently mutated in *BRCA1-*associated cancers and contributes to resistance to many genotoxic therapies [28–30].

Based on the mechanism of action, we find that these agents are genotoxic in nature, and the lesions caused by them are difficult to repair without intact *BRCA1* function. Previous literature reports have shown that *BRCA1*-mutant cells are highly sensitive to replicative DNA damage [14–16,34]. But unlike conventional chemotherapy drugs, the drug candidates identified by us showed preferential toxicity to mutant cells rather than normal mammary epithelial cells.

The NCI-60 correlation analysis provides orthogonal, database-derived support for treating these six compounds as a mechanistically related family rather than as five independent hits: the compounds selected based on initial experimentation, were confirmed to possess similar characteristics across in vitro cell lines.

The presence of various molecules with different kinetics of apoptosis also indicates different mechanisms. Differences between molecules might relate to types of lesions, replication requirement, and involvement of DNA damage checkpoints, as described for other genotoxic compounds [35–37]. This heterogeneity is important from a therapeutic point of view because fast acting molecules might act effectively together with DNA damage checkpoint inhibitors, while slow acting molecules might cooperate with replication stress inducers.

Resistance mechanisms for PARP inhibitors are often mediated through the restoration of HR or replication fork stabilization, through loss of *53BP1* or *BRCA1* reversion mutants [24–27]. If the DNA damage induced by these small molecules is resistant to partial restoration of DNA damage repair processes, then the use of these small molecules will still have utility in these settings. While the direct targets of these molecules are yet unknown, future work can assess their efficacy using preclinical models more reflective of clinical disease, such as patient-derived xenograft models, organoids, or *BRCA1*-mutant cells that restore HR function [24–27, 39–46].

The current study illustrates the usefulness of employing a genetically characterized human mammary epithelial cell system to screen for small molecules that specifically target DNA repair defects in *BRCA1*-mutant cells. The ability of these compounds to induce cell death in *TP53*-mutant and *TP53*-null cellular models supports their potential relevance for *BRCA1*-associated cancers, which frequently harbor *TP53* alterations. Overall, the current findings support continued investigation of these small molecules either alone or in drug combinations.

A search in CAS SciFinder Discovery Platform and Chemical Compound Database through the Chemical Abstracts Service (CAS), a division of the American Chemical Society in September 2026 retrieved the following information regarding the 5 *BRCA1* deficiency-selective compounds.

NSC667251 appeared in Patent US20250186411 A1, published June 12, 2025, entitled “Screening methods for identifying KLF15 transcription factor agonists and uses thereof for treating degenerative disorders associated with overactivation of IKKB, Patent US2015-62217986, application date 09-14-2015, entitled “Novel piperazine and piperidine derivatives, their synthesis and use as VDAC1 oligomerization inhibitors, Patent DE2011-102011118606, application date 11-15-2011, entitled “Preparation of stilbene compounds as PPARβ and PPARδ inhibitors useful in the treatment of PPARβ− and PPARδ-mediated diseases.” We have discussed earlier based on prior literature NSC636948 as an inhibitory ligand of peroxisome proliferator-activated β/δ (PPARβ/δ) and both NSC636948 and NSC667251 as inverse agonists of PPARβ/δ.

NSC636948 appeared in Patent JP05032606 A, published in 02-09-1993, entitled “Preparation of diarylacrylonitrile derivatives as nonlinear optical materials,” Patent JP61270747 A, published 12-01-1986, entitled “photopolymerizable compositions,” and Patent NL6512616, published 03-31-1966, entitled “Compounds with antiadrenal activity.”

NSC21683 appeared in Patent US20260049978 A1, published 02-19-2026, entitled “A method for treating CAG repeat expansion disorders using small molecules that selectively reduce expanded CAG transcript levels,” and Patent US2023-63606957, application date 12-06-2023, entitled “Screening methods for identifying KLF15 transcription factor agonists and uses thereof for treating degenerative disorders associated with overactivation of IKKB,”

NSC281383 and NSC136513 appeared in Patent US2012-61669213P, application date 07-09-2012, entitled “Bone morphogenetic protein pathway activation, compositions for ossification, and methods related thereto.” BMP pathway activation is growth promoting in cartilage and involves SMAD activation. Thus, further studies are required to understand how such activities may be connected to synthetic lethality with BRCA1-deficiency.

It is interesting that NSC667251 and NSC21683 appeared in patents referring to KLF15 transcription factors agonists that may be useful in disorders associated with overactivation of IKKB. These factors are relevant for kidney diseases as well as inflammatory and stress-induced fibrotic disorders. Further studies are warranted to explore connections between these pathways and synthetic lethality with BRCA1-deficiency. Further studies would be required to investigate connection between NSC667251, VDAC1 oligomerization inhibition, and synthetic lethality with BRCA1-deficiency, particularly since VDAC1 inhibition is predicted to inhibit apoptosis which is the opposite of what is observed in our studies.

Several limitations should be acknowledged. First, the molecular targets of the identified compounds remain unknown. Although all five compounds induced DNA damage signaling and apoptosis, additional studies will be required to determine their direct cellular targets and the precise nature of the DNA lesions generated. Second, the present study was performed primarily in cultured cell models and therefore does not address pharmacokinetics, toxicity, or in vivo antitumor efficacy. Third, although selective cytotoxicity was observed in *BRCA1*-deficient models, formal *BRCA1* complementation studies were not performed and will be required to definitively establish *BRCA1*-dependent sensitivity.

Future studies should focus on identification of the direct molecular targets of these compounds through chemo-proteomic and target-deconvolution approaches, assessment of efficacy in patient-derived organoids and xenograft models, and evaluation of activity in models of acquired PARP inhibitor resistance. Such studies will determine whether these compounds possess advantages over currently available DNA damage-targeting therapies.

In this study, we have identified five small molecule chemicals that can induce DNA damage as well as apoptosis in mammary epithelial and breast cancer cells with *BRCA1* deficiency. By employing a genetically defined human cell system, we have shown that loss of *BRCA1* creates a therapeutically exploitable vulnerability that may be targeted through selective induction of DNA damage and apoptosis. Importantly, the capability of these chemicals to induce apoptosis in the absence of wild-type *TP53* supports further preclinical evaluation in *BRCA1*-deficient tumors, including those harboring *TP53* mutations.

## Author Contributions

A.N. performed wet-bench experiments (primary and secondary drug screening, cell viability assays, and western blots) that are described when the El-Deiry Lab was at Penn State University. D.T.D. performed the flow cytometric analysis in the lab while at Penn State University. S.Z. assisted in setting up plates and providing the compounds for the chemical library screen and J-N.G. assisted in performing screening in the El-Deiry Lab at Penn State University. C.E-K. performed in silico analysis of chemical structures in publicly available databases for known activities and fingerprints. D.R. focused on the recognition of NSC636948 as a potential PPARβ/δ-directed ligand based on the published literature. K.S. assisted D.R. with Kaplan-Meier analysis of PPARD. J.G. and S.S.P.D. assisted the other authors in writing parts of the manuscript. W.S.E-D. conceptualized the project, designed the original drug screen, directed all aspects of the research, writing, editing of the manuscript, and financial support of the research. Using a personal subscription, W.S.E-D. performed the search in the CAS SciFinder Discovery Platform and Chemical Compound Database through the Chemical Abstracts Service (CAS), a division of the American Chemical Society in September 2026 to retrieve information regarding the 5 *BRCA1* deficiency-selective compounds.

## Acknowledgements

W.S.E-D. is an American Cancer Society Professor and is supported by the Mencoff Family University Professorship at Brown University. This work was supported at its early stages in part by an NIH grant (CA094975) to W.S.E-D. The NCI R01 grant support for the BRCA1-related research in the El-Deiry Lab ended in 2007. This work was submitted in November 2012 for presentation at the 2013 annual AACR meeting (A.N., Kim Nguyen, Bora Lim, David T. Dicker and W.S.E-D. “Drug screening for small molecules targeting BRCA1-deficient cells”). However, due to funding limitations, additional planned experiments were not performed, the poster was never presented at the 2013 AACR meeting, and the paper was never previously published.

